# Impaired methyl recycling induces substantial shifts in sulfur utilization in Arabidopsis

**DOI:** 10.1101/2025.03.09.642221

**Authors:** Benjamin J.M. Tremblay, Saeer A. Adeel, Maye Saechao, Yihan Dong, Eric Andrianasolo, John M. Steele, Annika Traa, Nilanth Yogadasan, Ishari Waduwara-Jayabahu, Barbara A. Katzenback, Rüdiger Hell, Markus Wirtz, Barbara A. Moffatt

**Affiliations:** Department of Biology, University Of Waterloo, Waterloo, Ontario, Canada N2L 3G1; Institut de Biologie Moléculaire des Plantes, UPR2357 du CNRS, Université de Strasbourg, Strasbourg, 67084, France; Centre for Organismal Studies (COS), Heidelberg University, 69120 Heidelberg, Germany

## Abstract

The tight coordination of sulfur metabolism and growth regulation is predicated upon nutrient availability. Central to this balancing act is the utilization of cysteine (Cys) for the formation of methionine (Met) and S-adenosylmethionine (SAM). Plants that are severely deficient in regenerating Met due to reduced methylthioadenosine (MTA) nucleosidase activity experience numerous developmental abnormalities. Here, we assess the developmental, metabolic, and regulatory effects associated with decreased MTA recycling. We show that MTA over-accumulation predominantly occurs in reproductive tissues and leads to reduced levels of Cys, Met, and SAM, as well as elevated S-adenosylhomocysteine. These disruptions of primary sulfur utilization also lead to the misregulation of energy metabolism and altered cell cycle progression. RNA-seq experiments show a general down-regulation of many developmental and reproductive genes. Targeted metabolite analyses demonstrate clear impacts on the methyl index which are reflected in the results of bisulfite-sequencing experiments including global alterations in CG gene-body methylation levels and decreases of CHG and CHH methylation in transposable elements. Our findings demonstrate the broad impacts of MTA metabolism on plant development, sulfur utilization and the maintenance of the methyl index.

## Introduction

Plants continuously monitor their nutrient status and environmental conditions to balance their growth and stress responses. Sulfur utilization is a crucial aspect of this balancing act. Plants fine-tune the flux of available sulfate towards various sulfur-containing compounds to coordinate growth and nutrient limitations with cysteine (Cys) biosynthesis being at the center of this regulation (Takahashi et al., 2011; Watanabe et al., 2021). Cys is a source of not only sulfur but also carbon and nitrogen precursors (Takahashi et al., 2011). Depending on developmental and environmental cues Cys utilization determines the flow of sulfur towards de novo biosynthesis of glucosinolates (GLS), glutathione (GSH) or methionine (Met) (Speiser et al., 2018).

Met limits the synthesis *S*-adenosylmethionine (SAM), which is the primary methyl donor for diverse methylation reactions, including DNA methylation. A substantial sink for SAM is the synthesis of metabolites including the hormone ethylene (Eth), the polyamines spermine (Spm), spermidine (Spd), and thermospermine (Tspm) and the metal ion chelator nicotinamide (NA; Ravanel et al., 1998). Using SAM to synthesize these key metabolites creates methylthioadenosine (MTA) as a by-product (Bürstenbinder et al., 2010; Waduwara-Jayabahu et al., 2012). The sulfur in MTA is recovered into Met in a group of reactions known as the Yang cycle (Ravanel et al., 1998). The enzyme MTA nucleosidase (MTN) catalyzes the first in this cycle (Ravanel et al., 1998). All *Arabidopsis thaliana* (Arabidopsis) Met salvage pathway mutants except for MTN exhibit no visible phenotypes and have been considered non-essential (Bürstenbinder et al., 2007; Friedman et al., 2011; Zierer et al., 2016).

Earlier analysis of MTN-deficient mutants showed accumulation of the MTA by-product to be detrimental in Arabidopsis. MTN activity is encoded by two functionally redundant genes in Arabidopsis: *MTN1* (*AT4G38800*) and *MTN2* (*AT4G34840*) (Bürstenbinder et al., 2010). The *mtn1-1mtn2-1* double mutant (with 14% residual MTN activity) accumulates increased levels of MTA even under normal growth conditions, leading to several developmental abnormalities, including delayed bolting, thickened vasculature, as well as both male and female sterility (Waduwara-Jayabahu et al., 2012). The basis of these phenotypes remains unknown, though previous studies have suggested that MTA-generating reactions are sensitive to MTA feedback inhibition (Herbik, 1997; Hyodo & Tanaka, 1986; Waduwara-Jayabahu et al., 2012). However, of these processes, only NA biosynthesis was decreased in *mtn1-1mtn2-1* plants grown under normal conditions. In fact, MTN-deficient mutant seedlings display wildtype-like Eth and Spd content, questioning the hypothesis that MTA feedback inhibition is the dominant reason for the substantial growth defects of *mtn* mutants (Waduwara-Jayabahu et al., 2012; Washington et al., 2016).

Here, we report our investigations of the biochemical basis for the severe developmental abnormalities associated with MTA accumulation, focussing on the sulfur assimilation pathway, and more specifically. the SAM cycle. Our results show that as MTA accumulates to high levels, a simultaneous reduction in SAM content occurs, indicating the existence of a possible homeostatic mechanism balancing sulfur utilization with MTA accumulation. This decline in sulfur assimilation leads to large-scale transcriptional changes and a loss in methylation potential and DNA hypomethylation, resulting in transposon reactivation.

## Materials and Methods

### Plant materials and growth conditions

*Arabidopsis thaliana* (Col-0) seeds were surface sterilized using chlorine gas and stratified in the dark at 4 °C for 72 h before sowing on half-strength MS medium with 0.8% agarose and 1% (w/v) sucrose (Murashige & Skoog, 1962). Seedlings were grown in a tissue culture chamber at 22 °C with continuous 100 µmol m-2 s-1 photosynthetically active radiation (PAR) for 14 d. Plants to be grown to maturity were transplanted to soil in individual pots and moved to a growth chamber with 150 µmol m^-2^ s^-1^ PAR in long day conditions. MTA feeding experiments were conducted as described previously (Bürstenbinder et al., 2010), sowing seeds on medium with either 500 µM MgSO_4_ or MTA (Abcam: ab141257) in concentrations ranging from 10 µM to 750 µM as a sulfur source and grown for 7 d. Polyamine supplemented media was prepared as previously described (Waduwara-Jayabahu et al., 2012)). Sampling was performed by flash freezing seedlings or unopened buds in liquid nitrogen before grinding to a powder and storing at -80 °C. The following T-DNA insertion lines were obtained from the Arabidopsis Biological Resource Centre (Ohio State University, USA): SALK_085385 (*mtn1-1*), SALK_071127 (*mtn2-1*), and SALK_022510 (*mtn2-5*). The *mtn1-1mtn2-1* and *mtn1-1mtn2-5* lines were generated and described previously (Waduwara-jayabahu, 2011; Waduwara-Jayabahu et al., 2012). PCR genotyping was performed as described previously (Waduwara-Jayabahu et al., 2012) using the primers listed in Supplementary Table 1.

### Plant phenotyping experiments

Vertically grown seedlings were tracked for 14 d by photographing the back of the petri dishes using a Nikon D3000 camera. The images were then used to measure the primary root length on ImageJ. Root lengths were standardized using an image of a ruler taken at the same focal length. The transition of a plant in soil from vegetative to reproductive development was marked by the visual emergence of the apical floral buds. The bolting time was measured from time of seed sowing to this milestone. At bolting, the number of rosette leaves was counted for each individual plant. Once the plant reached complete senescence, the height of the plant was measured from the base of the primary inflorescence stem to the top, in addition to the total number of siliques and bud clusters. Seeds produced by the plant were carefully harvested throughout its development. To determine the total number of seeds per plant, 500 seeds were initially weighed for each genotype and growth medium. The weight per 500 seeds was then used to quantify the total seed count for each replicate after weighing. The percent viability of the seeds was subsequently tested on half-strength MS medium.

### HPLC and quantification of metabolites

Metabolites were extracted from approximately 100 mg of frozen ground seedling or bud tissue powder using 1 mL 0.1 M HCl and clarified by centrifugation at 16,000 ⨉ *g* for 5 min at 4 °C. Extracts were stored at -80 °C. To measure thiols, 200 µL of 1/10th water-diluted extract was first reduced by adding 20 µL 1 M Tris (pH 8.3), 10 µL 10 mM DTT, and 25 µL 0.08 M NaOH before incubating in the dark for 1 h at room temperature. Derivatization was performed by addition of 25 µL 10 mM monobromobimane in acetonitrile and incubating in the dark for 15 min, before halting via addition of 705 µL 5% acetic acid. Separation was performed using HPLC on a Nova-Pak C18 (4.6 mm x 250 mm) column and signal detection with a Jasco 920 Intelligent Fluorescent Detector at 380 nm excitation and 480 emission. To measure adenosines, 300 µL of extract was derivatized by addition to 620 µL 620 mM citric acid-1-hydrate, 760 mM di-sodium hydrogen phosphate di-hydrate (pH 4) and 80 µL 45% chloroacetaldehyde before incubating at 80 °C for 10 min and subsequently cooled to room temperature on ice. Separation was performed as described earlier and signal detection using 280 nm emission with 410 excitation. For amino acid and polyamine measurements, 5 µL of extract was added to 35 µL 0.2 M boric acid (pH 8.8) and derivatized by addition of 20 µL 3 mg mL^-1^ AccQ Taq in acetonitrile and incubating at room temperature for 5 min, followed by incubating at 55 °C for 10 min, and finally diluting 5-fold with water. Separation was performed using HPLC on a Acquity BEH C18 column (150 mm x 2.1 mm) in an Acquity H-class UPLC and signal detection with an Acquity FLR-Detector set to 250 nm excitation and 395 emission. For NA measurements, 25 µL of extract was added to 75 µL 0.5 M borate buffer with 5 mM EDTA (pH 7.7) and derivatized by addition to 50 µL 12 mM 9-fluorenylmethyl chloroformate 9-fluorenylmothoxycarbonyl chloride in acetone before incubating for 45s at room temperature. The reaction was deactivated by addition of 50 µL 40 mM adamantan-1-amine amantadine hydrochloride in 75% acetone. Separation was performed using HPLC on a Phenomenex C18 column (4 mm x 3 mm) and signal detection with a Jasco 920 Intelligent Fluorescent Detector at 263 nm excitation and 313 nm emission. Quantification calculations from detection signals were performed using the software Empower Pro. For sulfate measurements, extracts were diluted 10-fold and separated in a Dionex ICS-1000 pump/detector with an Ion-PAC AS 9HC column (2 mm x 250 mm). Quantification calculations from detection signals were performed using the software Chromeleon (Dionex).

### Mass spec and data analysis

Unopened bud samples were extracted with 1 mL 0.1 M HCl and clarified by centrifugation at 16,000 ⨉ *g* for 5 min at 4 °C. Extracts were diluted with deionised water to make 1 mg/mL samples, then diluted to 1 µg/mL with 50% acetonitrile with 0.1% formic acid. Using a Velos LTQ Orbitrap Pro (ThermoFisher Scientific), positive and negative electrospray ionization were scanned in an ion trap (low resolution) with scan range 135-1000.

### TOR activity assays

For immunological analysis of TOR downstream phosphorylation, total soluble proteins were extracted from frozen 25 mg of unopened buds with a glass bead homogenizer and resuspended in homemade 2x Laemmli buffer at a 1:5 ratio of plant weight (mg) to buffer volume (µL). Samples were vortexed and boiled at 95°C for 5 min then transferred to ice. Samples were subsequently separated on a 12% 8M urea SDS-PAGE gel, and were transferred onto a polyvinylidene fluoride membrane. Detection of primary antibodies, S6K-p (Abcam; ab207399; 1:5 000) and S6K1/2 (Agrisera; AS12-1855; 1:5 000), was accomplished using a horseradish peroxidase-conjugated secondary antibody (1:20 000).

### Flow cytometry

Developmentally matched rosette leaves four and five were harvested from 14d old plants that were grown horizontally on half-strength MS medium or half-strength MS medium supplemented with 100 µM Spd, where seeds were sown at a density of 100 seeds per plate. Approximately 20 mg of leaf tissue was chopped finely using a new double-sided razor blade on a pre-chilled watch glass with 600 µL of freshly made ice-cold Aru buffer (Yang et al., 2019; 9.65 mL MgSO_4_ buffer (10 mM MgSO_4_·7H_2_O, 50 mM KCl, 5 mM HEPES), 1 mM DTT, and 0.1% (v/v) Triton X-100). The resulting extract was transferred to a 1 mL syringe with a homemade adapter to hold a 40 µm Nitex nylon mesh. The extract was filtered through this adapter into a 5 mL round-bottom polystyrene tube. The suspended nuclei were treated with RNase (50 µg/mL; BioShop, Toronto, Canada) for 10 min on ice and stained with propidium iodide (50 µg/mL). Samples were run on a BD Accuri C6 flow cytometer, operated at a flow rate of 14 µL/minute with acquisition settings as per (Galbraith & Lambert, 2012). Data analysis was performed using FlowJo software (BD), gating on individual ploidy peaks. The endoreduplication index was calculated by summing the products of the fraction of events at each ploidy level and the corresponding ploidy level value (i.e., 2C, 4C, 8C, 16C, 32C, or 64C). 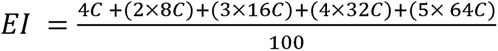

### MTN activity assay

MTN protein activity assays were performed as described by (Bürstenbinder et al., 2010). Briefly, 20 µg of total protein from unopened buds extracts in 50 mM potassium phosphate buffer (pH 7) were used in a xanthine oxidase-coupled enzyme assay, measuring absorbance at 470 nm every 10 min for 1 h. Enzyme activities were calculated from the rate of change of absorbance.

### RNA-seq experiments and analysis

RNA was extracted from approximately 50 mg samples of pooled frozen ground 14 d seedling or unopened bud tissue using the RNeasy mini Kit (Qiagen). RNA was DNase I-treated with the RNase-free DNase Set (Qiagen). Libraries were prepared and sequenced using either a HiSeq2500 or NovaSeq 6000 platform (Illumina) with a PE150 sequencing strategy by Novogene (Beijing, China) or Sickkids (Toronto, Canada). Reads were quality filtered and adapter-trimmed using Trim Galore! (Babraham Bioinformatics) and mapped onto the TAIR10 release of the Arabidopsis thaliana genome (Lamesch et al., 2012) with STAR (Dobin et al., 2013). Gene-level quantification based on the Araport11 *Arabidopsis thaliana* genome annotations (Cheng et al., 2017) was performed using StringTie (Pertea et al., 2015). The edgeR R package was used for differential expression analysis (Robinson et al., 2010). Gene Ontology (GO) enrichment analyses were performed for Biological Process (BP) terms using the topGO R package.

### BS-seq experiments and analysis

DNA was extracted from approximately 50 mg samples of pooled frozen ground unopened bud tissue using the DNeasy Plant Mini Kit (Qiagen). Bisulfite treatment, library preparation and sequencing using a HiSeq X Ten platform (Illumina) with a PE150 sequencing strategy were performed by Novogene. Reads were quality filtered and adapter-trimmed using Trim Galore! (Babraham Bioinformatics). Read mapping on the TAIR10 release of the Arabidopsis thaliana genome (Lamesch et al., 2012), read de-duplication and cytosine methylation calling were performed using Bismark (Krueger & Andrews, 2011). The DMRcaller R package was used for differential methylation analyses (Catoni et al., 2018). Global quantitation plots were created from data obtained using homer (Heinz et al., 2010). GO enrichment analyses were performed for BP terms using the topGO R package.

## Results

### Epigenetically restored *mtn1-1mtn2-1* plants have improved fertility and more normal growth

We previously reported that the fertility of the *mtn1-1mtn2-1* mutant can be partially restored by germinating and growing seedlings for 14 days on medium supplemented with either nicotianamine or a polyamine (Spd, Spm, or Tspm) before transplantation to soil (Waduwara-Jayabahu et al., 2012). Although the basis for this effect was not clear, it provided a means to study the *mtn1-1mtn2-1* phenotype without requiring recovery of the homozygous double mutant from segregating F2 populations. It also offered potential insights into the distinct mechanisms underlying MTA-driven sterility. Initially, we repeated the feeding experiment while carefully monitoring each compound’s effects on seedling growth and fertility. All polyamines restored the reduced primary root length of the double mutant while also leading to a minor increase in viable seed, though this latter effect was only significant for Spd (Figure 1b, c). We observed that seed obtained from Spd-treated plants generated progeny with greatly improved fertility levels when germinated on Spd-supplemented medium, and by the third generation (G3) the plants were substantially more WT-like in appearance, bolting time and in terms of seed yields compared to untreated *mtn1-1mtn2-1* plants (Figure 1a, d, e; Supplementary Figure 1a, b).

**Figure 1:**
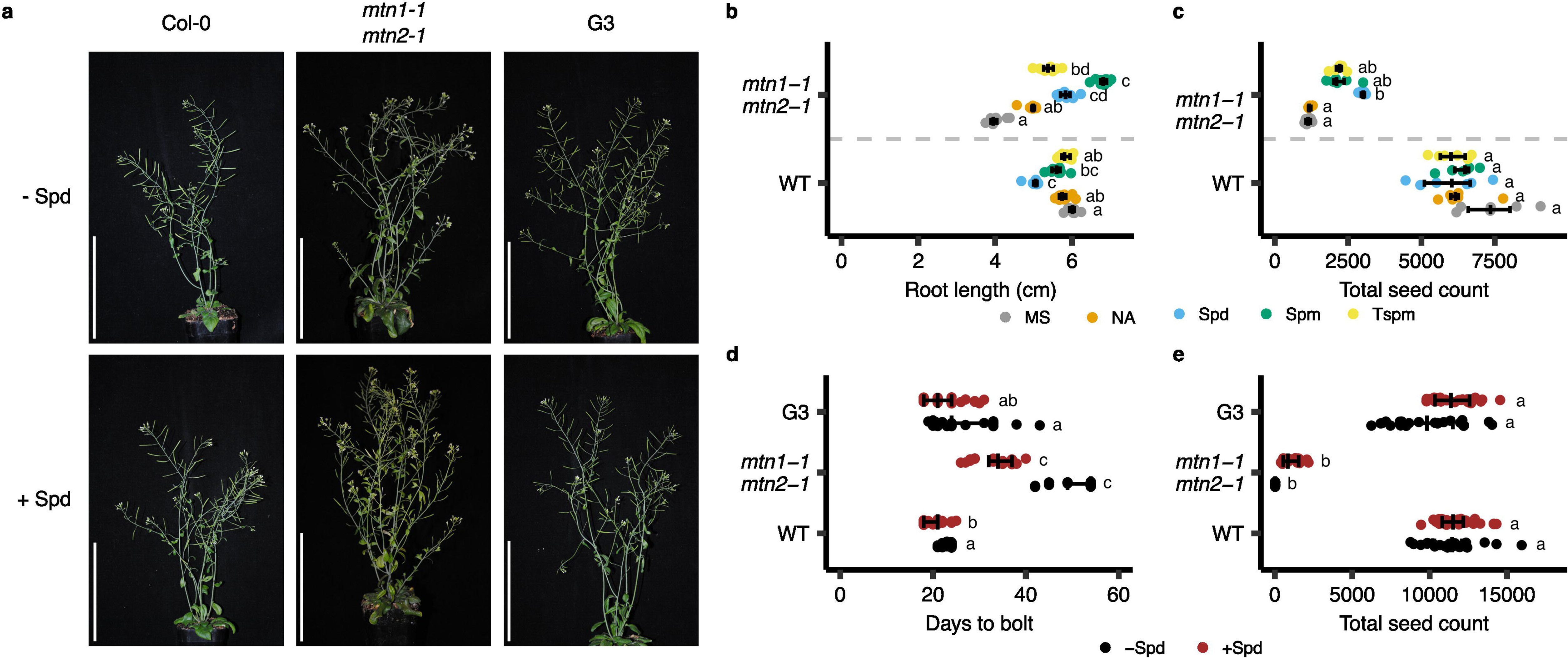
Spd supplementation partially restores *mtn1-1mtn2-1* growth. **a** Col-0 (WT), *mtn1-1mtn2-1*, and G3 plants were grown with or without Spd supplementation during the first two weeks of growth in petri plates. These plants were then transplanted into soil and representatives photographed upon silique ripening. Scale bar = 10 cm **b** Primary root lengths of WT and *mtn1-1mtn2-1* seedlings grown in petri plates for 14 d in the presence of NA, Spd, Spm or Tspm (n = 10). Seedlings were also grown in the absence of a treatment (MS). Statistical testing was performed using two-way Kruskal-Wallis tests with Dunn’s post-hoc test. Groups not sharing letters are significantly different (*P* < 0.05). **c** Plants exposed to the same treatment regimen from **b** were transplanted to soil and grown to term (n = 6). Seeds were collected at various stages of maturation and were allowed to dry. The total seed count per plant was assessed by initially weighing 500 seeds for each genotype and growth medium, with the weight per 500 seeds representing the seed yield of each replicate. Seed viability: WT (MS: 100%, NA: 100%, Spd: 100%, Spm: 99%, Tspm: 99%), *mtn1-1mtn2-1* (MS: 98%, NA: 96%, Spd: 100%, Spm: 98%, Tspm: 99%). **d** WT, *mtn1-1mtn2-1*, and G3 plants were grown as described for **a**, then transplanted to soil where the time till bolting was marked by the visual emergence of the apical floral buds (n = 17-31). Statistical testing was performed as described for **b**. **e** Plants from **d** were grown to term and the total seed count was quantified as described for **c** (n = 17-31). Seed viability: WT (MS: 100%, Spd: 99%), *mtn1-1mtn2-1* (MS: 98%, Spd: 99%), G3 (MS: 100%, Spd: 98%). Statistical testing was performed as described for **b**.

Given the broad range of phenotypic consequences of MTN deficiency at various developmental milestones (Waduwara-Jayabahu et al., 2012), we sought to assess the extent to which Spd supplementation restored normal development. The marked reduction in primary root length of *mtn1-1mtn2-1* seedlings compared to WT led us to examine their meristematic development, revealing smaller, and more disordered root apical meristems (Figure 2a). We found the increase in root length resulting from Spd treatment in *mtn1-1mtn2-1* seedlings (Figure 1b) coincided with a partial restoration of the meristematic zone, reaching approximately 1.5 times the length observed in *mtn1-1mtn2-1* seedlings grown on MS (Figure 2a, b). Notably, the loss of cellular organization within the meristematic zone of the *mtn1-1mtn2-1* mutant grown on MS was partially recovered following Spd treatment (Figure 2a). This cellular disorganization was observed not only in the quiescent centre (QC) but also extended to the columella cells and the epidermis, and cortex cell layers. In contrast, QC and columella disorganization was rarely seen in WT or G3 seedlings grown on either MS or Spd. G3 seedlings exhibited an overall more robust meristematic zone with sizes and organization similar to WT (Figure 2a).

**Figure 2:**
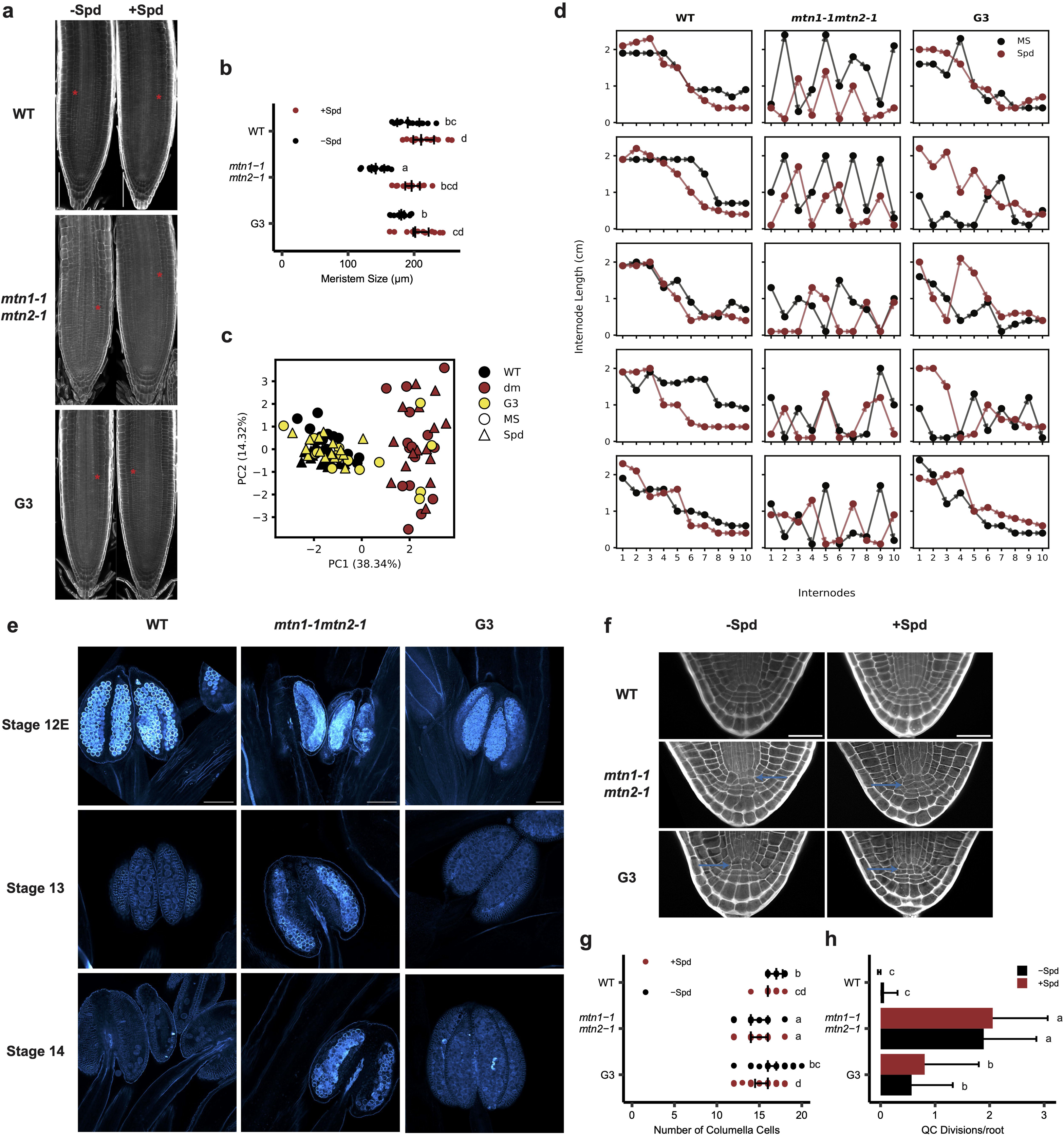
Partial restoration of *mtn1-1mtn2-1* phenotypes across developmental milestones. **a** Representative images of 7 day old Col-0 (WT), *mtn1-1mtn2-1*, and G3 roots obtained using the SR2200 cell wall stain. Scale bar = 100 µm. **b** The length of the meristematic zone was measured from isodiametric cells of the QC to the first cell that doubles in length in the cortex cell layer, indicated by the red asterisk in **a**. Statistical testing was performed using two-way Kruskal-Wallis tests with Dunn’s post-hoc test. Groups not sharing letters are significantly different (*P* < 0.05; n=15). **c** After WT, *mtn1-1mtn2-1* (dm), and G3 plants reached approximately 25 cm in height, the first ten internodes between siliques were measured on the primary inflorescence branch for both Spd treated and untreated plants and summarized in a PCA (n=15). **d** The distribution of internode lengths of five representative plants for WT, *mtn1-1mtn2-1*, and G3 and Spd treated or untreated depicted based on internode and its length. **e** Callose deposition in anthers of WT, *mtn1-1mtn2-1*, and G3 plants during stages 12E, 13, and 14 of floral development, stained with aniline blue. **f** WT (n=38, 47), *mtn1-1mtn2-1* (n=53, 37), and G3 (n=47, 51) seeds were imbibed on media with or with Spd supplementation for 24h and the cell walls of the mature embryo root were visualized by propidium iodide staining. Scale bar = 100 µm. **g** Both the number of columella cells and the number of cell divisions (blue arrow) in the QC were quantified. Statistical testing was performed as in **b**.

The improvements in bolting time and seed yield in Spd-treated *mtn1-1mtn2-1* plants prompted us to further investigate the phenotypic effects of Spd on floral development and anther maturation. We measured the first ten internode lengths of the primary inflorescence branch, and found the patterns of *mtn1-1mtn2-1* internode lengths were distinct from those of WT and G3 plants based on a principle component analysis (PCA), with some variability in the internodes of G3 plants germinated without Spd clustering near *mtn1-1mtn2-1* plants (Figure 2c). Overall, *mtn1-1mtn2-1* plants exhibited poorly organized phyllotaxy compared to WT. The inconsistency of lengths from one internode to another is consistent with previous observations of their shortened plant stature, abnormal plant architecture, and fasciation. The impairment of proper internode development due to MTN deficiency was only partially restored in G3 plants, with Spd-treated G3 plants showing more consistent internode lengths as seen in WT (Figure 2d). In addition to the abnormal phyllotaxy of the *mtn1-1mtn2-1* mutant, we assessed the level of callose deposition in *mtn1-1mtn2-1* anthers during flower stages 12E to 14, as callose deposition and degradation are fundamental anther development milestones (Lu et al., 2014; Sanders et al., 1999). Anthers of *mtn1-1mtn2-1* flowers contained clumpy and deformed pollen, and flowers at stages 13 and 14 did not dehisce (i.e. release pollen grains) normally (Figure 2e). After three generations of growth on Spd-supplemented media, G3 plants showed a significant reduction in callose deposition compared to *mtn1-1mtn2-1*, along with the increased release of pollen grains in stage 14 flowers (Figure 2e).

Given the severe defects observed in seed production and root apical meristem physiology in *mtn1-1mtn2-1*, we further explored whether MTN deficiency also disrupts embryogenesis in addition to impairing floral meristem development. To evaluate the embryonic development of *mtn1-1mtn2-1* plants, we quantified columella cell numbers and QC divisions of mature *mtn1-1mtn2-1* embryos, as these cell types are established during embryogenesis and provide key developmental insights (Forzani et al., 2014). Mutant *mtn1-1mtn2-1* embryos had significantly fewer columella cells than WT, or G3 plants grown on media in the absence of Spd supplementation (-Spd), while their QC cells were twice as likely to divide, and exhibit abnormal asymmetrical divisions (Figure 2f, g, h). In addition, both *mtn1-1mtn2-1* and G3 embryos exhibited irregularities in the cell organization of the root meristem (Figure 2f). Although Spd treatment of the mother plants (as seedlings for 14 d) did not induce noticeable effects, G3 embryos were nevertheless similar to the WT in terms of their cell organization. Since multiple aspects of plant development are partially restored in G3 plants, we saw this transgenerationally stable restoration as an opportunity to examine metabolic and regulatory changes associated with MTA over-accumulation.

### Moderately MTN-deficient *mtn1-1mtn2-5* mutant plants exhibit mild developmental abnormalities and improved fertility

To investigate how varying levels of MTN deficiency impact plant development, we aimed to generate an allelic series of MTN-deficient mutants. For that reason, we generated a moderate MTN-deficient mutant, *mtn1-1mtn2-5*, by crossing the *mtn1-1* single mutant with a new MTN2 allele, *mtn2-5* (SALK_022510). This double mutant retained 28% residual MTN activity, twice the amount present in the severe *mtn1-1mtn2-1* double mutant (Figure 3a). Phenotypic analysis of *mtn1-1mtn2-5* revealed that its primary root growth was less inhibited by MTA-supplemented medium compared to *mtn1-1mtn2-1*, indicating a clear dose-dependent response to MTA accumulation (Figure 3b). Notably, the additional residual MTN activity in *mtn1-1mtn2-5* substantially mitigated the negative effects of MTA accumulation, restoring much of the primary root growth and significantly reducing delays in bolting observed in the severe mutant (Figure 3b-e). Despite displaying an increased number of bud clusters, indicative of floral meristem activation commonly associated with MTN deficiency, *mtn1-1mtn2-5* remained fertile. Remarkably, it produced more than twice the seed yield of WT plants, contrasting sharply with the sterility of *mtn1-1mtn2-1* (Figures 3f-g). This moderate MTN-deficient mutant demonstrates that even modest changes in MTN activity can significantly affect growth and sterility.

**Figure 3:**
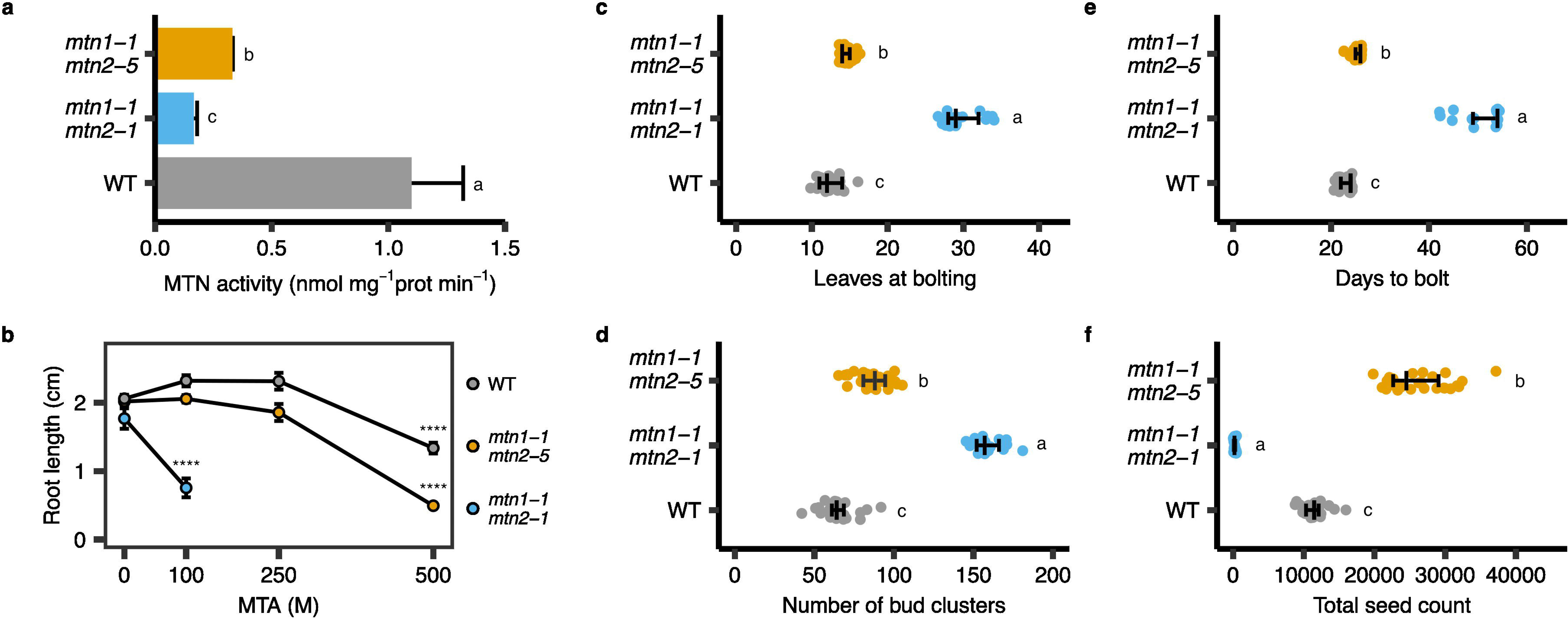
The *mtn1-1mtn2-5* mutant displays an intermediate phenotype. **a** MTN activity assays performed using bud clusters from WT, *mtn1-1mtn2-1*, and *mtn1-1mtn2-5* plants (n = 3). Data from WT and *mtn1-1mtn2-1* plants have been previously published. All samples are from the same experiment. Statistical testing was performed using two-sided Student’s t-tests with Holm correction for multiple testing. Groups not sharing letters are significantly different (*P* < 0.05). **b** Primary root lengths of Col-0 (WT), *mtn1-1mtn2-1*, and *mtn1-1mtn2-5* plants grown on plates containing MTA-supplemented medium (n = 8-64). Statistical testing was performed using two-way Kruskal-Wallis tests with Dunn’s post-hoc test. Only comparisons to Col-0 are shown (\*\*\*\**P*<0.0001). Col-0 and *mtn1-1mtn2-1* data from this assay have been previously published by Waduwara-Jayabahu et al., 2012. **c-f** Phenotyping data of Col-0 (WT), *mtn1-1mtn2-1*, and *mtn1-1mtn2-5* plants (n = 17-32), including the number of leaves at bolting, number of days until bolting, the number of bud clusters at senescence, and total seed count upon harvest, respectively. Statistical testing was performed using two-way Kruskal-Wallis tests with Dunn’s post-hoc test. Groups not sharing letters are significantly different (*P* < 0.05).

### Increased MTA accumulation in buds leads to reduced fertility

As a starting point in our characterization of the metabolic effects of MTA accumulation, we performed targeted analysis of 29 metabolites, including key compounds related to the SAM cycle and primary sulfur assimilation in seedling and bud tissues of our mutants, with and without Spd supplementation (Figure 4a; Supplementary Data 1). These measurements revealed that while the *mtn1-1mtn2-1* mutant seedlings accumulate two-fold higher levels of MTA as compared to WT, the buds of mature plants contain more than ten-fold higher levels of MTA (Figure 4c, c). This led us to hypothesize that reproductive tissues have a greater demand for metabolites whose syntheses generate MTA as a by-product and thus are more dependent on MTN activity. Steady state NA levels were lower in buds and seedlings of MTN-deficient mutants (Figure 4d-e). However, Spd was not affected in any of the MTN mutants (Figure 4f; Supplementary Figure 2a). These findings suggest that the biosynthesis of the two metabolites are under differential regulatory control and respond differently to changes in the availability of their precursor compounds. In buds, this may be in part due to increased accumulation of the Spd precursor putrescine (Put) (Supplementary Figure b-c).

**Figure 4:**
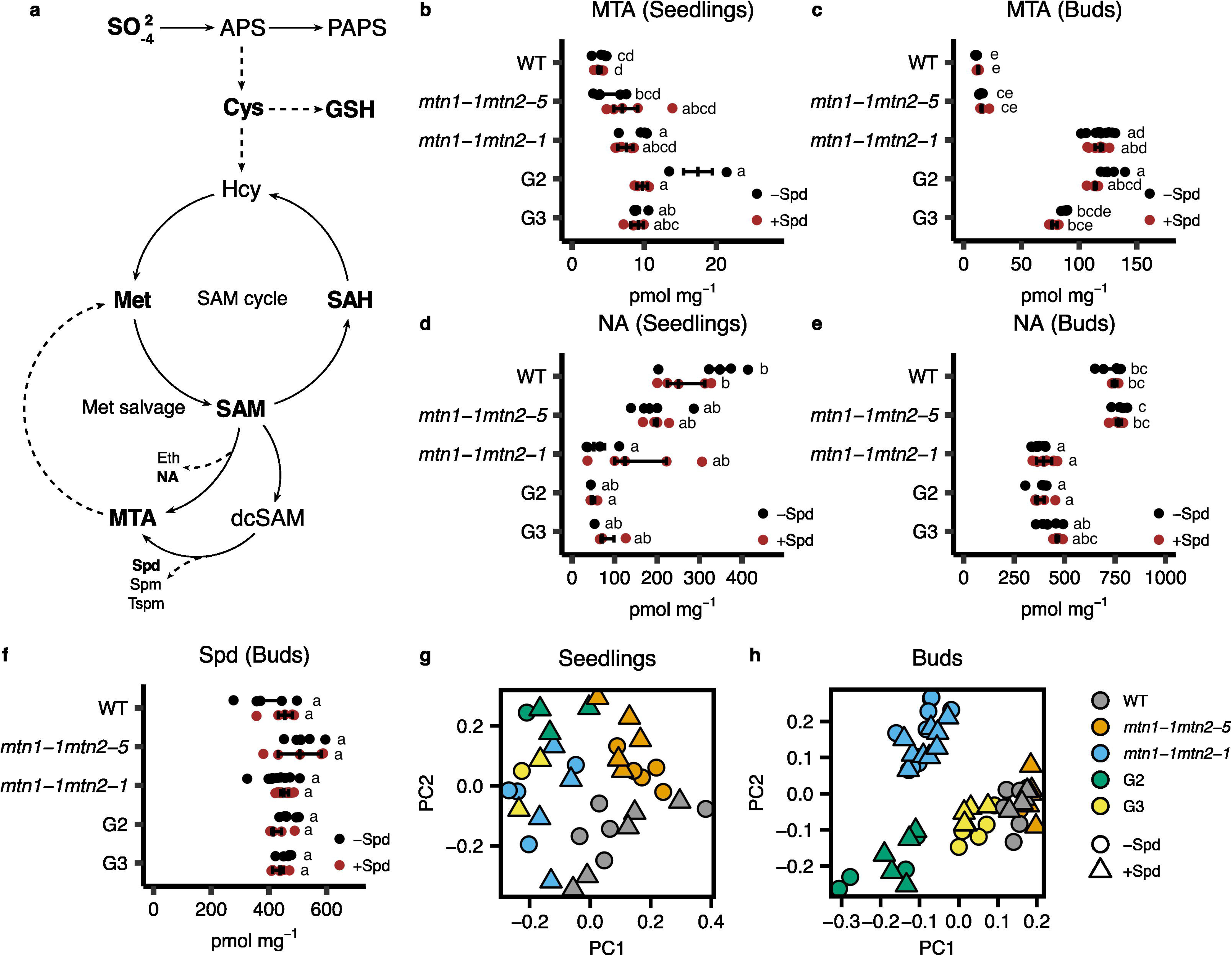
MTA accumulation mis-regulates primary sulphur assimilation. **a** Schematic of the primary sulphur assimilation pathway, including the SAM cycle and Met salvage. Dashed lines hide intermediate steps. Bolded metabolites have been quantified in this study. **b-hf** Individual metabolite quantification data including MTA in seedlings, MTA in buds, NA in seedlings, NA in buds, and Spd in buds, respectively (n = 4-5). Statistical testing was performed using two-way Kruskal-Wallis tests with Dunn’s post-hoc test. Groups not sharing letters are significantly different (*P* < 0.05). **g, h** PCA of all metabolite quantification data from seedlings and buds in this study, respectively, including Col-0 (WT), *mtn1-1mtn2-1*, *mtn1-1mtn2-5*, G2, and G3 plants. Seedlings were grown in the presence or absence of Spd.

The substantially stronger increase in MTA accumulation in mutant buds compared to seedlings was accompanied with an increased global impact on all measured metabolite levels resulting in greater separation in the principal component analyses between the mutant and WT samples as compared to seedlings (Figure 4b, c). Interestingly, despite the moderate *mtn1-1mtn2-5* mutants exhibiting developmental abnormalities, the metabolite profile of these plants clustered very close to the WT samples in the PCA of buds (Figure 4c). In contrast to *mtn1-1mtn2-1*, the *mtn1-1mtn2-5* mutants did not accumulate significantly higher levels of MTA in buds, likely because the additional residual MTN activity is sufficient to prevent MTA accumulation (Figure 4d-e). Since the most significant developmental difference between the *mtn1-1mtn2-5* and *mtn1-1mtn2-1* mutants was their fertility, we concluded that the reduced fertility of *mtn1-1mtn2-1* is primarily caused by MTA accumulation in the bud. Indeed, we even observed a slight decrease in MTA accumulation in the more fertile G3 buds compared to *mtn1-1mtn2-1* buds (Figure 4e).

Additionally, we observed during the process of sampling unopened buds from *mtn1-1mtn2-1* branches often led to subtending flower buds becoming fertile (Supplementary Figure 3). This observation suggests that individual flowers likely do not accumulate sufficient MTA to cause sterility. Instead, excess MTA accumulated in the bud cluster is likely diffused or transported in a basal direction, maintaining the opened flowers in a state of sterility.

### Primary sulfur assimilation metabolites steady-state levels are affected by MTA

We next examined the impact of MTA accumulation and MTN deficiency on the SAM cycle (Figure 4a). Despite being grown in sulfur-sufficient conditions, *mtn1-1mtn2-1* buds contained significantly decreased levels of Met and SAM (Figure 5a, b). Previous work examining the levels of these metabolites in mutants of other biosynthetic enzymes of the Met salvage pathway revealed that this pathway is only necessary to replenish levels of Met and SAM in sulfur-insufficient conditions, leading us to conclude that MTA itself may be acting in some unknown manner to disrupt regulation of the SAM cycle independent of sulfur availability (Zierer et al., 2016). This was further evidenced by elevated levels of SAH and reduced levels of Cys in the mutant, despite no observable differences in the levels of free sulfate (Figure 5c-e). Levels of SAM, SAH and Cys were restored in G3 buds, but Met was not (Figure 5a-c, e). In combination with the lack of significant differences for these metabolites in *mtn1-1mtn2-5* buds, these results confirmed the importance of SAM cycle metabolites in maintaining fertility. In particular, SAH accumulation in buds correlated well with MTA accumulation and decreased fertility.

**Figure 5:**
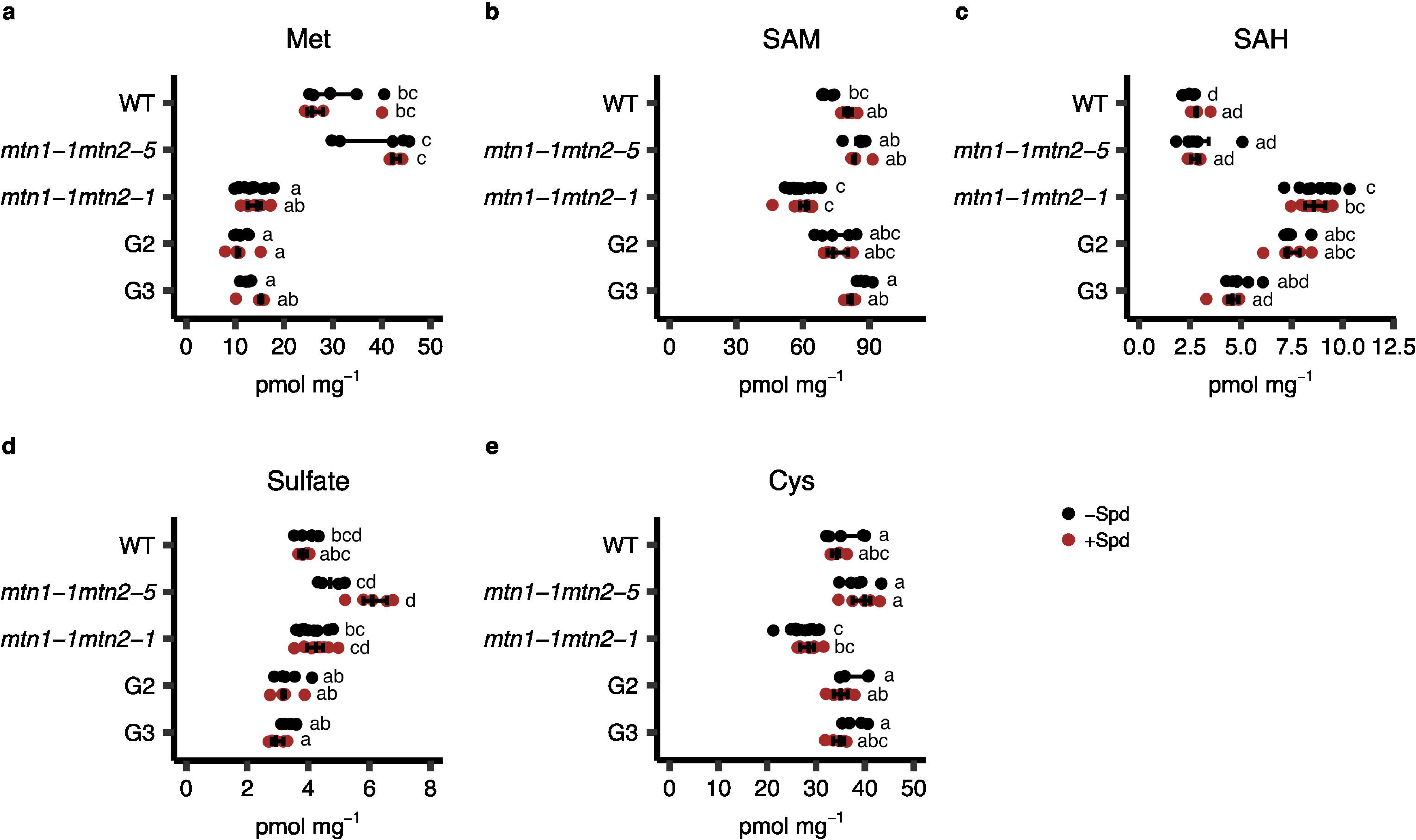
Effects of MTA accumulation on SAM cycle metabolites is independent of sulphate levels. **a-e** Individual metabolite quantification data of Met, SAM, SAH, sulphate, and Cys, respectively, from buds of Col-0 (WT), *mtn1-1mtn2-1*, *mtn1-1mtn2-5*, G2, and G3 plants having been grown in the presence or absence of Spd supplementation (n = 4-5). Statistical testing was performed using two-way Kruskal-Wallis tests with Dunn’s post-hoc test. Groups not sharing letters are significantly different (*P* < 0.05).

### Altered energy metabolism and cell cycle changes are likely secondary effects of MTA

Since the Yang cycle recycles the sulfur moiety in MTA, we investigated the effects of the MTN-deficient phenotype on sulfur sensing mechanisms in plants. Previous work has shown that plants sense sulfur deficiency and reduce their growth via the target of rapamycin (TOR) pathway, which acts as a master regulator of energy metabolism (Dong et al., 2022). We first examined the levels of the amino acids which we had measured in our metabolite measurement experiments, as regulation of TOR function requires proper homeostasis of free amino acid levels (Kim & Guan, 2019). There were slight increases in the abundances of most of the measured amino acids in *mtn1-1mtn2-1* seedlings, which were generally unchanged in G3 seedlings (Supplementary Figure 4a). In buds, there was a wider range of both increased and decreased levels of amino acids in *mtn1-1mtn2-1* plants. Reductions of the levels of Ile, Leu, Lys, and Tyr were reversed in G3 buds, indicating MTA-dependent mis-regulation of amino acid metabolism in *mtn1-1mtn2-1* plants.

We next investigated whether MTA accumulation in *mtn1-1mtn2-1* plants led to an impairment in the cell cycle and meristem activity, which are under tight regulation of the TOR signaling network (Pfeiffer et al., 2016). We observed early induction of the endocycle in *mtn1-1mtn2-1* mutants (Supplementary Figure 4b, d). These plants had an endoreduplication index approximately 1.5-fold higher than WT plants, alongside a substantial pool of 8C and 16C nuclei, which was largely recovered in G3 plants. Finally, we measured levels of the phosphorylated and unphosphorylated forms of S6K, the downstream target of TOR kinase activity (Wullschleger et al., 2006). Although we could detect reduced average levels of S6K (both phosphorylated and unphosphorylated) in *mtn1-1mtn2-1* buds, as well as a reduced ratio of phosphorylated to unphosphorylated S6K, these changes were not significantly different from WT (Supplementary Figure 4c; Supplementary Figure 5). The lack of clear evidence for disruption of TOR activity in a manner linked to MTA accumulation led us to conclude that while MTN deficiency is likely associated with alterations in energy metabolism and cell cycle progression, these changes may be secondary to other effects of MTA accumulation.

### Differential expression of developmental genes in *mtn1-1mtn2-1* buds

To provide further evidence for our hypothesis that MTA accumulation is a main driver of the observed developmental differences in MTN-deficient mutants, we next compared their transcriptomes. Since *mtn1-1mtn2-1* and *mtn1-1mtn2-5* seedlings did not accumulate MTA, we expected to find only minor changes in their transcriptomes. Indeed, the transcriptomes of the mutant and WT seedlings were highly correlated, with few differentially expressed genes between *mtn1-1mtn2-1* and WT (Figure 6a; Supplementary Figure 6; Supplementary Dataset 2). We then proceeded with an RNA-seq experiment using buds from *mtn1-1mtn2-1* mutants, which do accumulate high levels of MTA, as well as from the partially recovered G3 plants. This time we observed larger differences in the overall transcriptome of *mtn1-1mtn2-1* mutants as compared to WT (Figure 6b; Supplementary Figure 7; Supplementary Dataset 3). The majority of differentially expressed genes in *mtn1-1mtn2-1* buds were strongly down-regulated. A gene ontology enrichment analysis revealed many of these genes were involved in cell wall organization, extracellular region, and pollination among others (Figure 6c). Spd treatment was specifically further associated in the differential expression of genes regulating the response to abiotic stimulus and hypoxia. A large proportion of the differentially expressed genes in *mtn1-1mtn2-1* buds had the opposite differential expression pattern when comparing G3 and *mtn1-1mtn2-1* buds, suggesting an association between these specific pathways and the lack of fertility of the mutant (Figure 6d-e).

**Figure 6:**
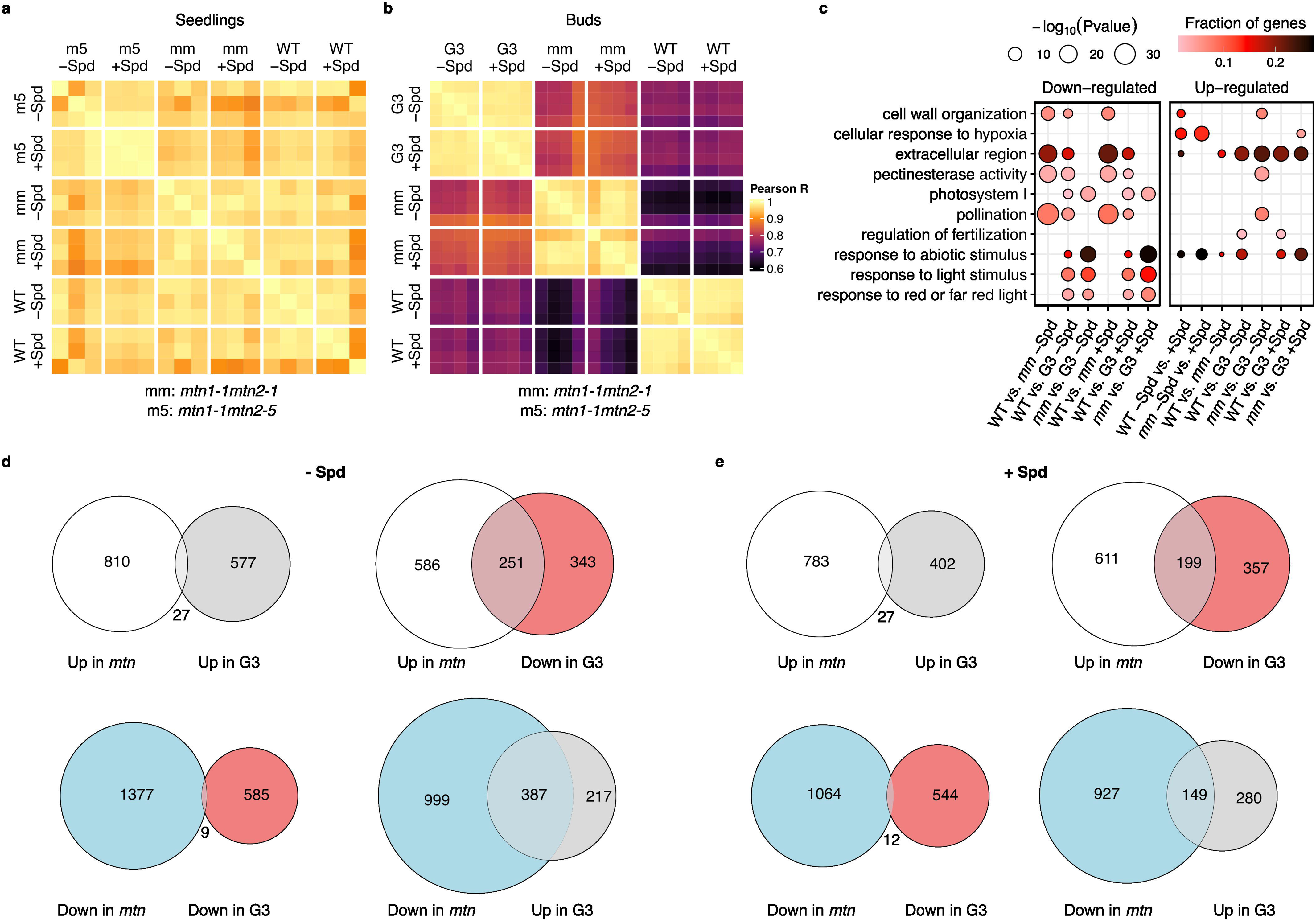
Major transcriptional changes underpin the *mtn1-1mtn2-1* phenotype. **a** Pearson correlation coefficient (*R*) matrix from RNA-seq transcript quantification of Col-0 (WT), *mtn1-1mtn2-1* (mm) and *mtn1-1mtn2-5* (m5) seedlings with or without Spd supplementation. **b** Same as **a** for RNA-seq transcript quantification of WT (Col-0), *mtn1-1mtn2-1* (mm), and G3 buds grown with or without Spd supplementation. **c** Enriched GO terms from up- and down-regulated gene sets between WT, *mtn1-1mtn2-1* (mm), and G3 comparisons of differentially expressed genes. The size of the circle represents the magnitude of the -log_10_ of the P-value, and the colour represents the fraction of genes in each gene set associated with the enriched GO term. **d-e** Venn diagrams of the number of overlapping differentially expressed genes between *mtn1-1mtn2-1* (*mtn*) plants versus WT (Col-0) and G3 plants versus *mtn1-1mtn2-1*, with or without Spd supplementation.

### Abnormal distribution of CG methylation and reduced levels of CHG and CHH methylation lead to gene silencing and transposable element hypomethylation

The differential gene expression analysis also revealed that many genes annotated as “transposable_element_gene” in Araport11 (Cheng et al., 2017) were up-regulated in *mtn1-1mtn2-1* buds (Figure 7a). We hypothesized the up-regulation of transposable element genes could be due to their hypomethylation, as they are normally silenced by DNA methylation. A loss of methylation potential was supported by a decreased SAM/SAH ratio in buds, also known as the methyl index, an indicator of methylation capacity (Figure 7b) (Caudill et al., 2001; Hoffman et al., 1980)). We proceeded with profiling genome-wide DNA methylation in buds using bisulfite-sequencing (BS-seq), which in addition to profiling the methylation status of transposable elements would allow us to determine whether any potentially heritable DNA methylation changes were associated with the epigenetic fertility restoration phenotype of G3 plants.

**Figure 7:**
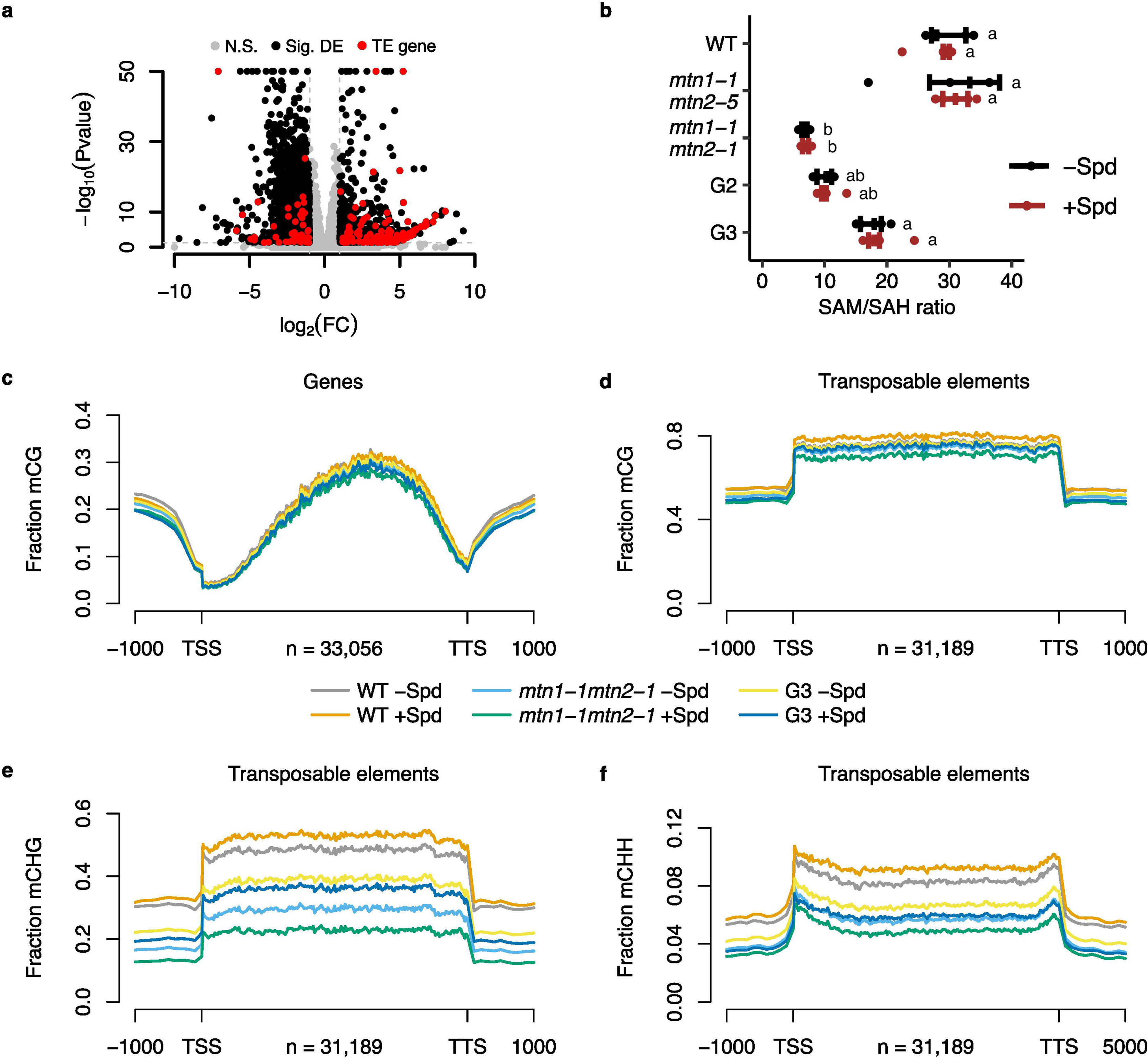
MTA accumulation is associated with major loss of CHG DNA methylation. **a** Volcano plot of the -log_10_ P-value and log_2_ fold-change (FC) of differentially expressed genes between WT and *mtn1-1mtn2-1* buds. Points representing significantly differentially expressed genes are coloured in black (as opposed to grey for non-significant), and significantly differentially expressed transposable element genes are coloured in red. **b** Ratio of SAM to SAH in buds as an indicator of the methyl index. Quantification data is from Col-0 (WT), *mtn1-1mtn2-1*, *mtn1-1mtn2-5*, G2, and G3 plants with or without Spd supplementation (n = 4-5). Statistical testing was performed using two-way Kruskal-Wallis tests with Dunn’s post-hoc test. Groups not sharing letters are significantly different (*P* < 0.05). **c, d** Metagene plots of average CG DNA methylation, including 1 Kbp upstream and downstream, for genes and transposable elements, respectively, from the Araport11 annotations. DNA methylation data is from BS-seq of WT, *mtn1-1mtn2-1*, and G3 buds, with or without Spd supplementation. **e, f** Metagene plots of average CHG and CHH DNA methylation, respectively, of transposable elements as for **d**.

Our analysis revealed thousands of differentially methylated regions (DMRs) in the CG context in *mtn1-1mtn2-1* and G3 buds compared to WT, both in terms of hypo and hypermethylation (Supplementary Figure 8a; Supplementary Dataset 4). Many of these CG DMRs were in common between *mtn1-1mtn2-1* and G3 buds, however we observed that 462 hyper-methylated DMRs in *mtn1-1mtn2-1* buds were in turn hypo-methylated in G3 buds. The presence of a large number of DMRs did not affect the average profile of methylated CGs over gene bodies or transposable elements, suggesting these events are likely not because of systematic changes in CG methylation capacity (Figure 7c-d). Indeed, many of these CG DMRs were specifically enriched in down-regulated genes in the RNA-seq related to cell and pollen growth GO terms (Supplementary Figure 9). Spd appeared to have little effect, with no detectable changes in WT methylation and relatively few DMRs in *mtn1-1mtn2-1* and G3 buds (Supplementary Figure 10a).

MTA accumulation appeared to instead have a much greater effect on global methylation levels in the CHG and CHH contexts. We observed a near 50% decrease in average CHG and CHH methylation in transposable elements in *mtn1-1mtn2-1* buds, with partial recovery in G3 buds (Figure 7e-f). There were 4957 hypo-methylated DMRs in the CHG context in *mtn1-1mtn2-1* buds, compared to only 5 hyper-methylated DMRs (Supplementary Figure 8b). G3 buds had far fewer CHG context DMRs, with 887 hypo-methylated (of which 457 were in common with *mtn1-1mtn2-1*) and 11 hyper-methylated DMRs. As before Spd appeared to have little effect within the same generation (Supplementary Figure 10b). Far fewer CHH context DMRs could be identified, likely as a result of methylation in this context being present in low amounts (Supplementary Figure 8c).

## Discussion

The Yang cycle plays an important role in allowing plants to replenish their pools of SAM without requiring additional sulfur inputs (Figure 4a) (Bürstenbinder et al., 2010; Zierer et al., 2016). SAM is an essential compound which is used as a methyl donor in many methylation reactions throughout cells and is a precursor for the biosynthesis of ethylene, nicotianamine and polyamines. While loss of a functional Yang cycle does not impair development of Arabidopsis under normal conditions, near total loss of MTN activity associated with overaccumulation of MTA leads to several developmental and reproductive deficiencies even under sulfur-sufficient conditions (Waduwara-Jayabahu et al., 2012; Zierer et al., 2016). Making use of the MTN-deficient mutant *mtn1-1mtn2-5* with moderate residual MTN activity (Figure 3), we confirmed the reproductive failure phenotype seen in severe *mtn1-1mtn2-1* mutants to be specific to their high levels of MTA (Figure 4). Curiously, we did not detect a significant change in MTA abundance in *mtn1-1mtn2-5* relative to WT, suggesting a tipping point exists in the ability of the plant to accommodate reduced MTN activity; this lies between 14% and 28% of the MTN activity in WT plants. Mutant *mtn1-1mtn2-1* 2^nd^ and 3^rd^ generation transgenerationally-restored plants (G2 and G3, respectively) by treatment with 14d Spd also had decreased levels of MTA in buds in association with their increase in fertility (Figure 1, Figure 2, Figure 4). Partial phenotypic rescue of root meristematic development, abnormal phyllotaxy, anther physiology, and embryonic development were also triggered with the initial 14d Spd treatment (Figure 2). The level of MTA accumulation required to impair reproductive organs is likely very close to the high levels seen in *mtn1-1mtn2-1*, as G3 buds are fertile despite having only a moderate decrease in MTA accumulation and the untreated double mutant produces occasional fertile siliques. Spd treatment also altered the steady-state levels of several metabolites in the primary sulfur assimilation pathways, such as increased SAH and reduced SAM and Cys, changes which were reversed in G3 plants (Figure 5). Overall, the data establish a clear link between high levels of MTA accumulation, misregulation of primary sulfur metabolism, and reproductive failure irrespective of the actual sulfur levels available to the plant.

Since we could not identify a clear link between the changes in primary sulfur metabolism and downstream phenotypic effects, we performed RNA-seq of MTN-deficient seedlings and buds. Our analyses revealed the biggest changes in the transcriptome of *mtn1-1mtn2-1* buds, which correlated with the high levels of MTA accumulation in that tissue (Figure 4e; Figure 6a-b). Differentially expressed genes were enriched in pathways related to fertilization, validating previous observations concerning male and female sterility and the disruption of anther development (Figure 6c, Figure 2e) (Waduwara-Jayabahu et al., 2012). Additionally, we noticed an up-regulation in the expression of TE genes in both *mtn1-1mtn2-1* seedlings and buds (Figure 5d-g). Crucially many fewer of these genes were expressed in *mtn1-1mtn2-5* or G3 samples, leading us to conclude the increased expression of these genes were possibly associated with the high levels of MTA accumulation and plant sterility. Increased expression of transposable element genes has been linked to stress responses and global DNA hypomethylation in plants (Hudson et al., 2011; Roquis et al., 2021). Our RNA-seq experiment showed clear signs of up-regulation of stress-related genes in *mtn1-1mtn2-1* plants. Furthermore, a previous report has demonstrated DNA hypomethylation in *hog1* mutant plants which lack the primary SAH hydrolase enzyme activity, leading to a disruption in the SAM cycle (Figure 4a) (Ouyang et al., 2012). The reduction in DNA methylation in *hog1* mutants is likely the result of the decrease in SAM (or an increase in SAH leading to feedback inhibition of methylation reactions), similar to what we observed in *mtn1-1mtn2-1* mutants (Figure 5a-c). BS-seq of *mtn1-1mtn2-1* buds revealed a near 50% decrease in CHG and CHH methylation at sites which are involved in small RNA-guided silencing of TE methylation (Figure 7e-f) (Van Ex et al., 2011). As expected, G3 buds had intermediate levels of mCHG and mCHH over TEs. This suggests the methylation potential of the plant is highly sensitive to varying levels of MTA, with its accumulation having a direct impact on availability for methylation.

Plants finely regulate sulfur metabolism to balance growth and stress responses, with Cys biosynthesis playing a central role in this coordination. Our study reveals that MTA accumulation disrupts this delicate balance, causing severe misregulation of primary sulfur utilization, resulting in reduced levels of Cys, Met and SAM, and impairing downstream processes such as DNA methylation and nicotianamine biosynthesis. The reduced utilization of sulfur for Cys biosynthesis suggests a shift in sulfur assimilation toward secondary metabolism, such as GLS biosynthesis, in MTN-deficient mutants. This is further supported by our detection of additional, highly abundant unknown compounds in *mtn1-1mtn2-1* samples from HPLC experiments, potentially linked to elevated secondary sulfur metabolites used for GLS biosynthesis (Supplementary Note 1). While the majority of GLS are converted into simple nitriles, a recent study suggested that sulfur is reallocated back to Cys production in a retrograde pathway under sulfur deprivation (Sugiyama et al., 2021). MTN-deficient plants may be utilizing this avenue for relieving the deprivation of primary sulfur metabolites.

Beyond its impact on sulfur assimilation, MTA accumulation influences various developmental pathways, underscoring its wide-reaching effects on plant growth, as evidenced by altered root meristems and impaired floral development. A plant’s ability to overcome the disruptions caused by MTA accumulation appears to play a critical role in determining its fertility, as seen in G3 mutants, where slightly lower MTA levels than in *mtn1-1mtn2-1* mutants correlate with partially restored fertility. In conclusion, these data demonstrate the essential role of MTN activity in plant sulfur utilization, independent of other enzymes in the Yang cycle, highlighting its critical contribution to growth and development.

## Availability of data and materials

All RNA-seq and BS-seq datasets generated in this study are available from the NCBI GEO repository under the accession GSE288809. Mutant lines generated in this study are available upon request to the corresponding author.

## Acknowledgements and Funding

We are grateful to Zachary T. Hull for performing MTN enzyme activity assays and the Metabolomics Core Technology Platform of the Excellence Cluster “CellNetworks” (University of Heidelberg, Germany) for amino acid and polyamine measurements. B.J.M.T. was funded in part by a Ontario/Baden-Wurttemberg (OBW) Summer Research Program scholarship from the Ontario Ministry of Training, Colleges, and Universities (Canada). Work in the Heidelberg lab was supported by funding (project no. 235736350 and 544882710) from the Deutsche Forschungsgemeinschaft (German Research Foundation, DFG) granted to R.H. and M.W..

## Author Contributions

B.J.M.T., S.A.A., and B.A.M. conceptualized the study and wrote the manuscript with input from all authors. B.J.M.T., S.A.A., M.W., R.H., I.W.-J., and B.A.M. designed the research. B.J.M.T., S.A.A., M.S., Y.D., E.A., J.M.S., A.T., B.A.K., N.Y., I.W.-J., and B.A.M. performed the research. B.J.M.T., S.A.A., Y.D., E.A., and B.A.M. analyzed the data. B.A.M. supervised the study.

