## Supplementary Table 1 for "Impaired methyl recycling induces substantial shifts in sulfur utilization in Arabidopsis"

**Supplementary Table 1: Genotyping primers used in this study.**

| Primer name | Description | Sequence |
|---|---|---|
| mtn1-1F | Genotyping SALK_085385 with LBb1.3. | TGACGGAGACCAACTCCATAC |
| mtn1-1R | | GAGGCTCTTCCTTTGGTCAAC |
| mtn2-1F | Genotyping SALK_071127 with LBb1.3. | CCTTGCTTACGTGGCATAAAC |
| mtn2-1R | | GGAAAGGGCAAAAATATATGG |
| mtn2-5F | Genotyping SALK_022510 with LBb1.3. | ACTGTGCCAACACTCTCAACC |
| mtn2-5R | | AAGATTTCCGCTTCCTGAAAG |
| LBb1.3 | Used with SALK primers. | ATTTTGCCGATTTCGGAAC |
