## Supplementary Figure 1 for "Impaired methyl recycling induces substantial shifts in sulfur utilization in Arabidopsis"

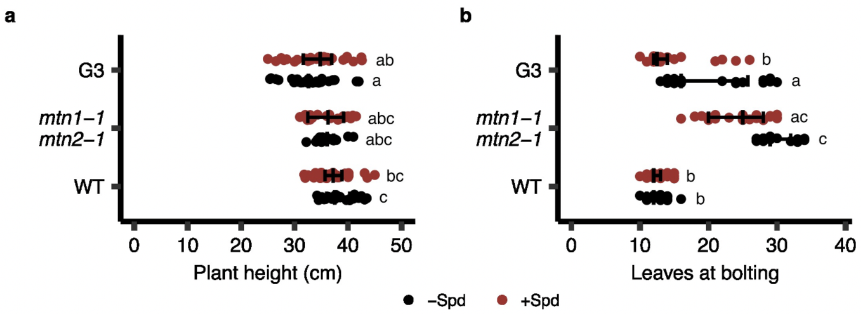


**Supplementary Figure 1: Further phenotypic analyses of the effect of Spd on *mtn1-1mtn2-1* growth.**

**a** Col-0, *mtn1-1mtn2-1*, and G3 plants were grown with or without spermidine (Spd) supplementation during the first two weeks of growth in petri plates (n = 17-31), and then transplanted into soil. The plant height was measured from the base of the primary inflorescence stem to the top when the apical bud cluster failed to produce new flowers. Statistical testing was performed using two-way Kruskal-Wallis tests with Dunn's post-hoc test. Groups not sharing letters are significantly different ($P < 0.05$). **b** The transition from vegetative growth to flowering was marked by the emergence of the floral bud cluster, at which point the rosette leaves were counted. Statistical testing was performed as per **a**.
