## Supplementary Figure 2 for "Impaired methyl recycling induces substantial shifts in sulfur utilization in Arabidopsis"

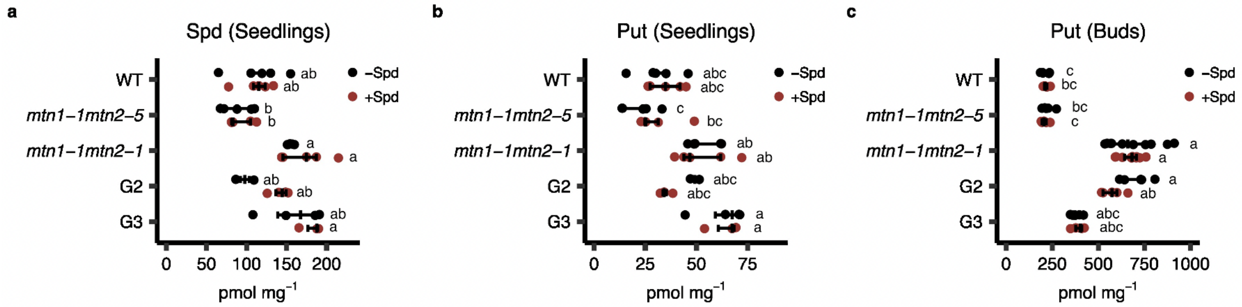


**Supplementary Figure 2: Polyamine levels are not reduced in MTN-deficient plants.**

**a-c** Metabolite quantification data of Spd in seedlings, Put seedlings, and Put in buds, respectively, from Col-0 (WT), *mtn1-1mtn2-1*, *mtn1-1mtn2-5*, G2, and G3 plants grown with or without Spd supplementation (n = 4-5). Statistical testing was performed using two-way Kruskal-Wallis tests with Dunn's post-hoc test. Groups not sharing letters are significantly different ($P < 0.05$).
