## Supplementary Figure 3 for "Impaired methyl recycling induces substantial shifts in sulfur utilization in Arabidopsis"

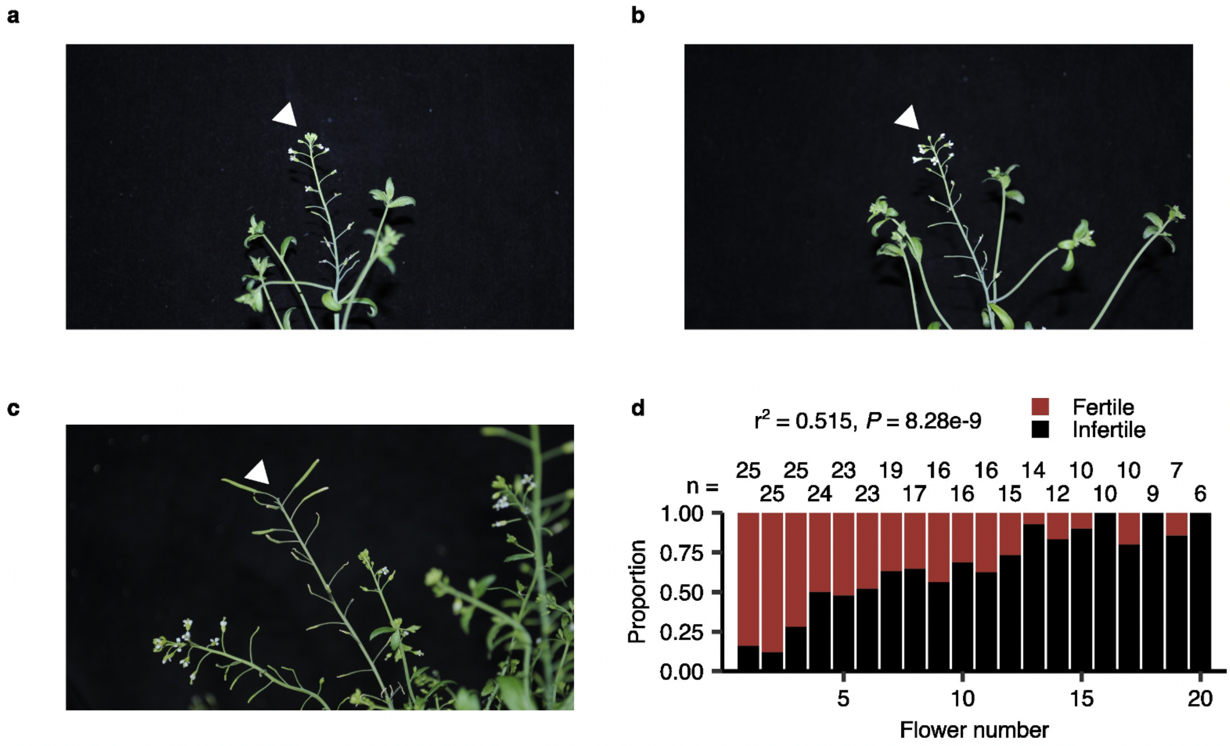


**Supplementary Figure 3: Unopened bud decapitation in *mtn1-1mtn2-1* restores fertility.**

**a-c** Photographs of a representative *mtn1-1mtn2-1* shoot before, immediately after, and 2 wk after decapitation of the apical unopened bud cluster (white arrow), respectively. **d** Relative proportion of fertile and infertile siliques on branches with a decapitated apical unopened bud cluster (n = 25). The siliques were counted in ascending order of the flowers remaining after decapitation. The data was fitted to a linear model to test whether a significant relationship exists between the proportion of fertile flowers and the distance from the site of decapitation (i.e., flower number; adjusted R-squared = 0.515, *P*-value = 8.28e-9).
