## Supplementary Figure 4 for "Impaired methyl recycling induces substantial shifts in sulfur utilization in Arabidopsis"

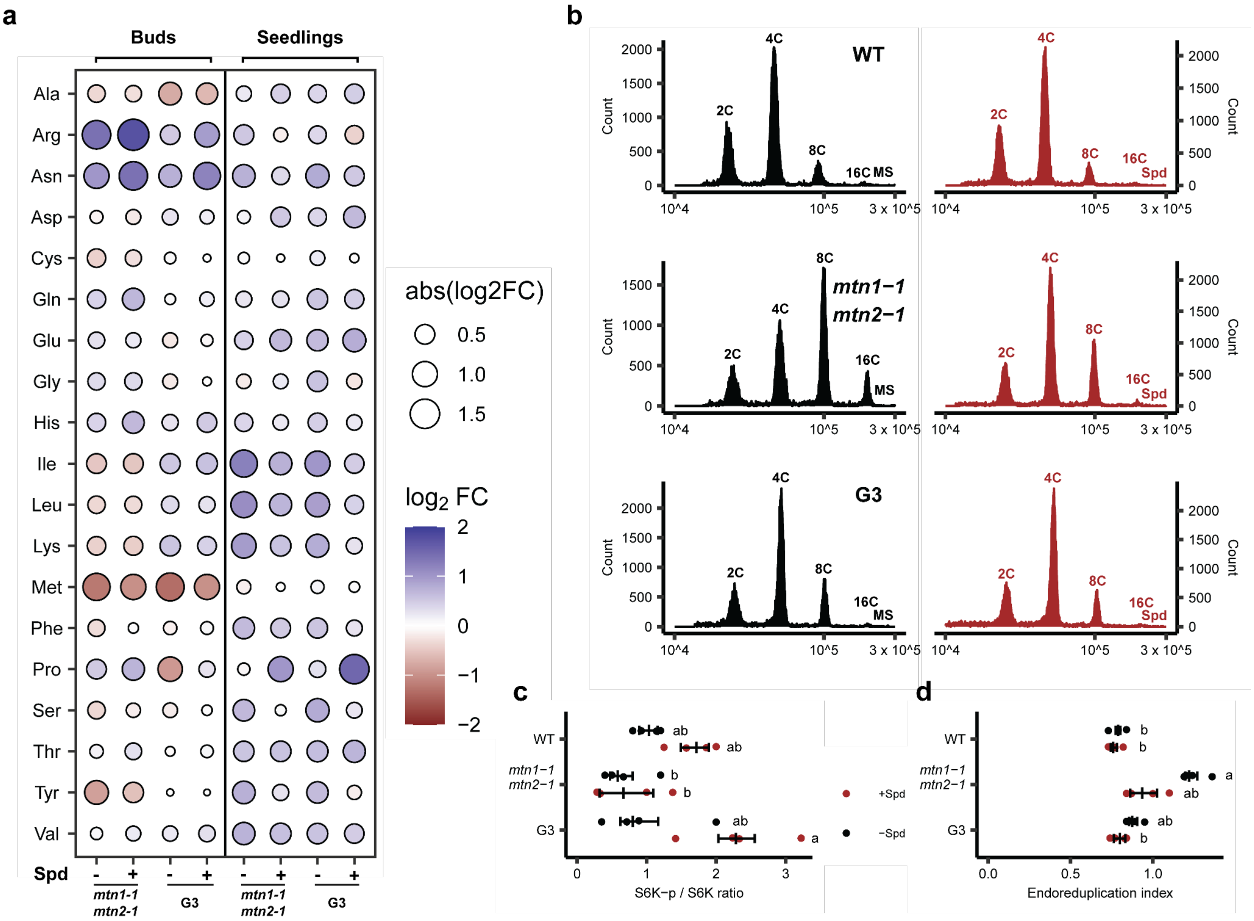


**Supplementary Figure 4: Lack of clear evidence of TOR mis-regulation by MTA.**

**a** Average fold change (FC) levels of 19 amino acids in *mtn1-1mtn2-1* and G3 seedlings and buds, with or without Spd supplementation, compared to Col-0 (WT; n = 4-5). The $\log_2$ FC levels are shown by colour, and the absolute magnitude shown by the size of the circle. **b** The cell ploidy of 14d old WT, *mtn1-1mtn2-1*, and G3 seedlings grown horizontally on MS (black-filled histogram plots) and Spd (red-filled histogram plots) media (n = 4) were measured using flow cytometry with developmentally matched rosette leaves. Four major peaks in these representative histograms are from cells with ploidy levels of 2C, 4C, 8C, and 16C, respectively. **c** Ratio of phosphorylated S6K (S6K-p) to unphosphorylated S6K (as a measure of TOR activity) in WT, *mtn1-1mtn2-1*, and G3 buds grown with or without Spd supplementation (n = 4). **d** The endoreduplication index was calculated from the percentage of isolated nuclei within each major peak. Statistical testing was performed using two-way Kruskal-Wallis tests with Dunn's post-hoc test. Groups not sharing letters are significantly different ($P < 0.05$; n=4). Statistical testing was performed as for **c**.
