## Supplementary Figure 5 for "Impaired methyl recycling induces substantial shifts in sulfur utilization in Arabidopsis"

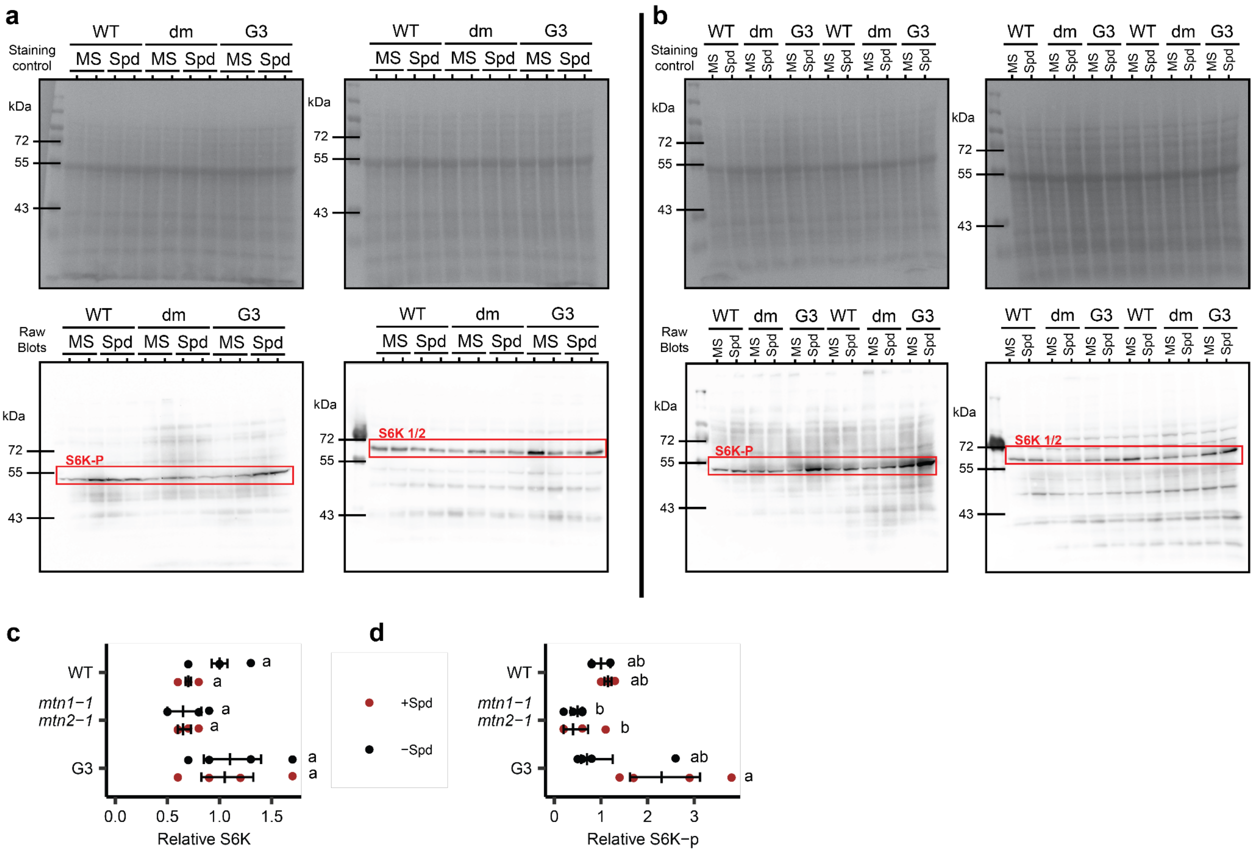


**Supplementary Figure 5: S6K levels are largely not affected in *mtn1-1mtn2-1*.**

**a-b** Raw, uncropped Coomassie Blue loading control blots and immunoblots showing unphosphorylated and phosphorylated S6K levels from Col-0 (WT), *mtn1-1mtn2-1* (dm), and G3 buds grown with or without Spd supplementation. Biological replicates 1 and 2 are shown in **a**, while replicates 3 and 4 are shown in **b**. **c-d** Relative quantification of unphosphorylated and phosphorylated S6K levels from blots shown in **a-b**, respectively, (n = 4). Statistical testing was performed using two-way Kruskal-Wallis tests with Dunn's post-hoc test. Groups not sharing letters are significantly different ($P < 0.05$). Protein levels are shown relative to the average levels in Col-0 buds grown without Spd supplementation.
