## Supplementary Figure 6 for "Impaired methyl recycling induces substantial shifts in sulfur utilization in Arabidopsis"

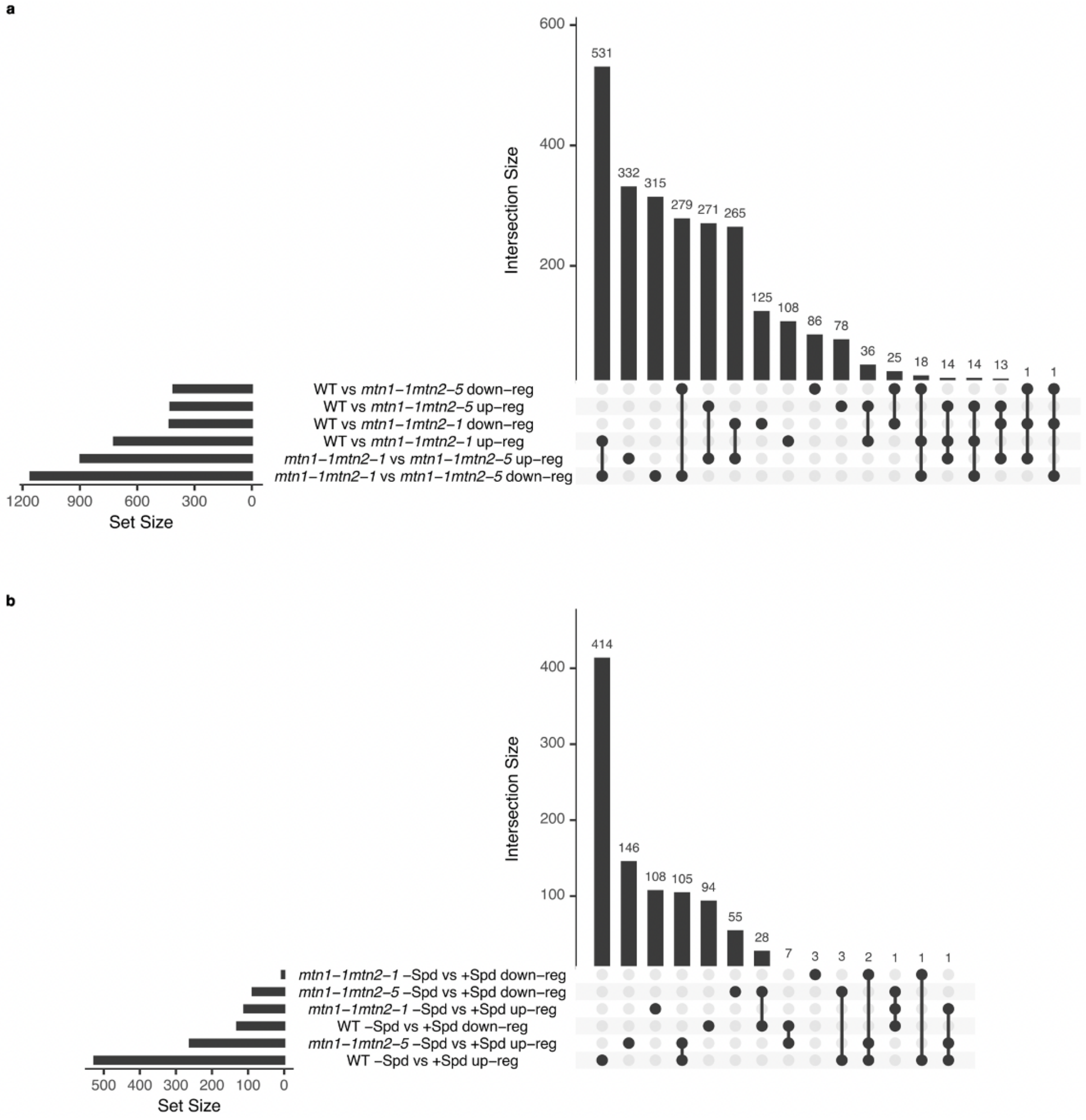


**Supplementary Figure 6: Transcriptome changes in *mtn1-1mtn2-1* seedlings are minor.**

**a** UpSet plot of overlapping gene sets in comparisons between RNA-seq of Col-0 (WT), *mtn1-1mtn2-1*, and *mtn1-1mtn2-5* seedlings without Spd supplementation. **b** UpSet plot of overlapping gene sets from comparisons between treatments (with and without Spd supplementation) of samples from **a**.
