## Supplementary Figure 7 for "Impaired methyl recycling induces substantial shifts in sulfur utilization in Arabidopsis"

a

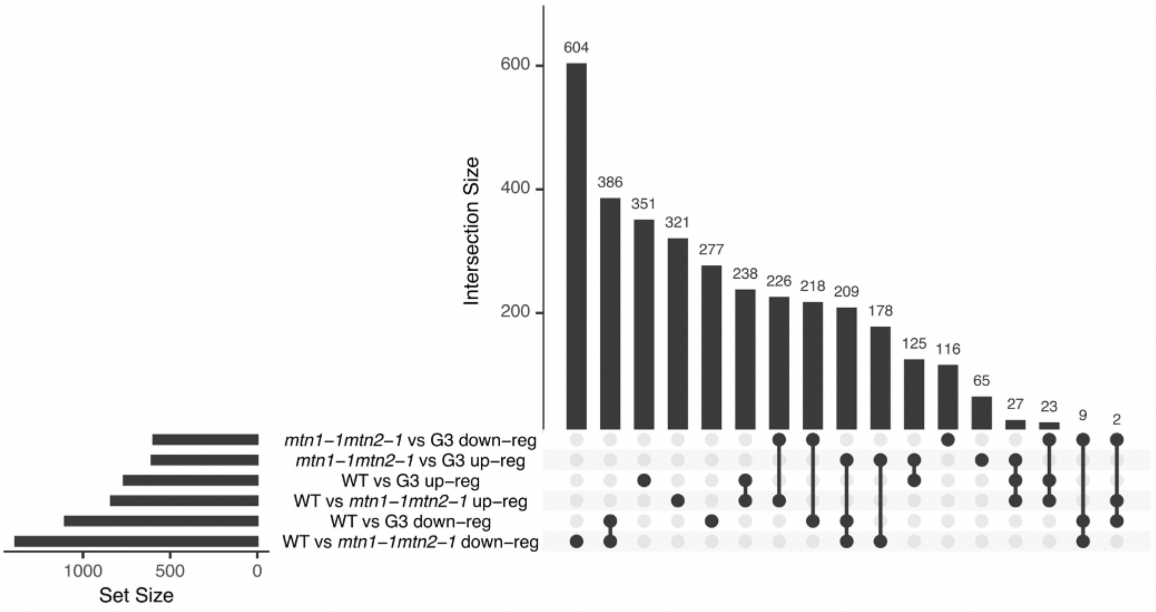


b

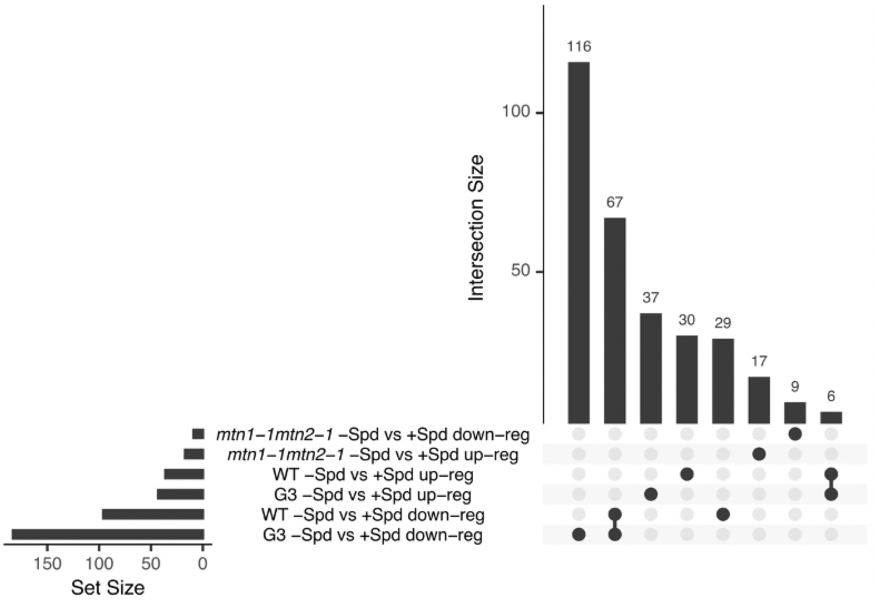


**Supplementary Figure 7: The transcriptome of *mtn1-1mtn2-1* buds diverges strongly from Col-0 buds.**

**a** UpSet plot of overlapping gene sets in comparisons between RNA-seq of Col-0 (WT), *mtn1-1mtn2-1*, and *mtn1-1mtn2-5* buds without Spd supplementation. **b** UpSet plot of overlapping gene sets from comparisons between treatments (with and without Spd supplementation) of samples from **a**.
