## Supplementary Figure 8 for "Impaired methyl recycling induces substantial shifts in sulfur utilization in Arabidopsis"

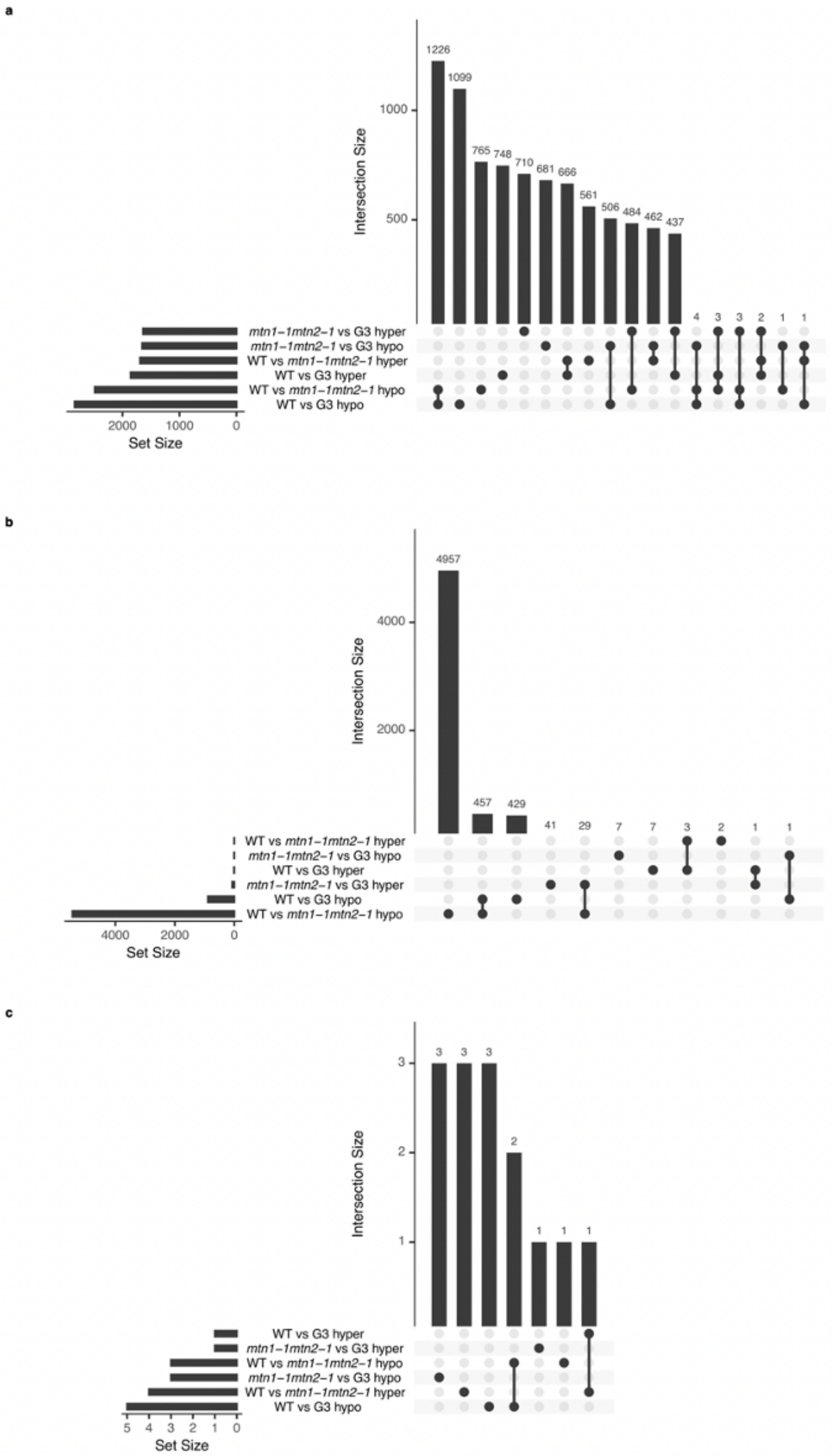


**Supplementary Figure 8: Global changes in DNA methylation in *mtn1-1mtn2-1* and G3.**

**a** UpSet plot of overlapping sets of mCG DMRs in comparisons between BS-seq of Col-0 (WT), *mtn1-1mtn2-1*, and G3 buds without Spd supplementation. **b** Same as **a** in the mCHG context. **c** Same as **a** in the mCHH context.
