## Supplementary Figure 9 for "Impaired methyl recycling induces substantial shifts in sulfur utilization in Arabidopsis"

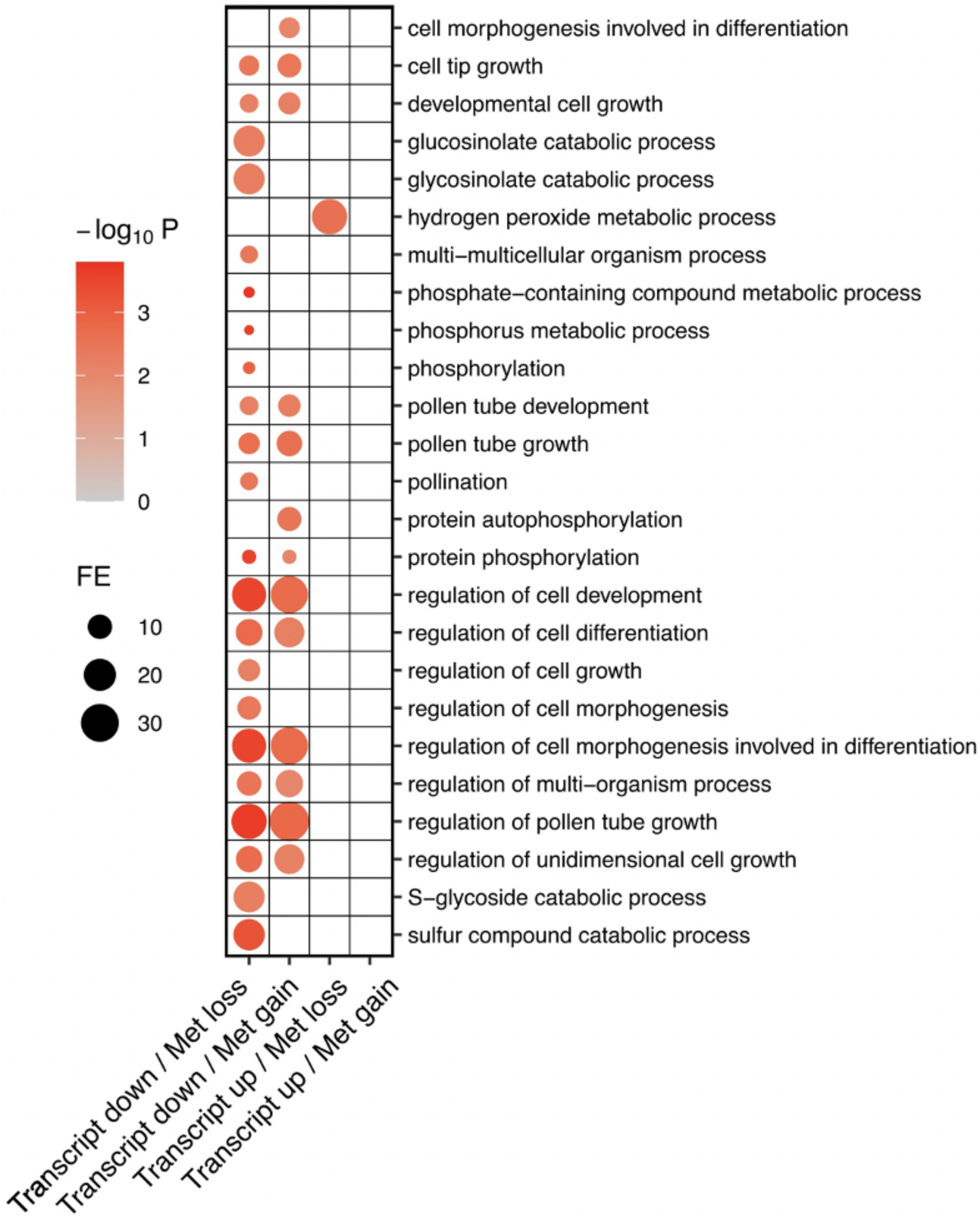


**Supplementary Figure 9: Changes in DNA methylation in *mtn1-1mtn2-1* lead to changes in transcript abundance of many pollen genes.**

GO enrichment of significantly differentially expressed genes in RNA-seq of *mtn1-1mtn2-1* buds with associated mCG DMRs. The colour of the circle represents the magnitude of the $-\log_{10}$ of the P-value, and the size of the circle represents the fold enrichment of genes with the associated GO term in the gene set over expected background counts.
