## Supplementary Figure 10 for "Impaired methyl recycling induces substantial shifts in sulfur utilization in Arabidopsis"

**a**

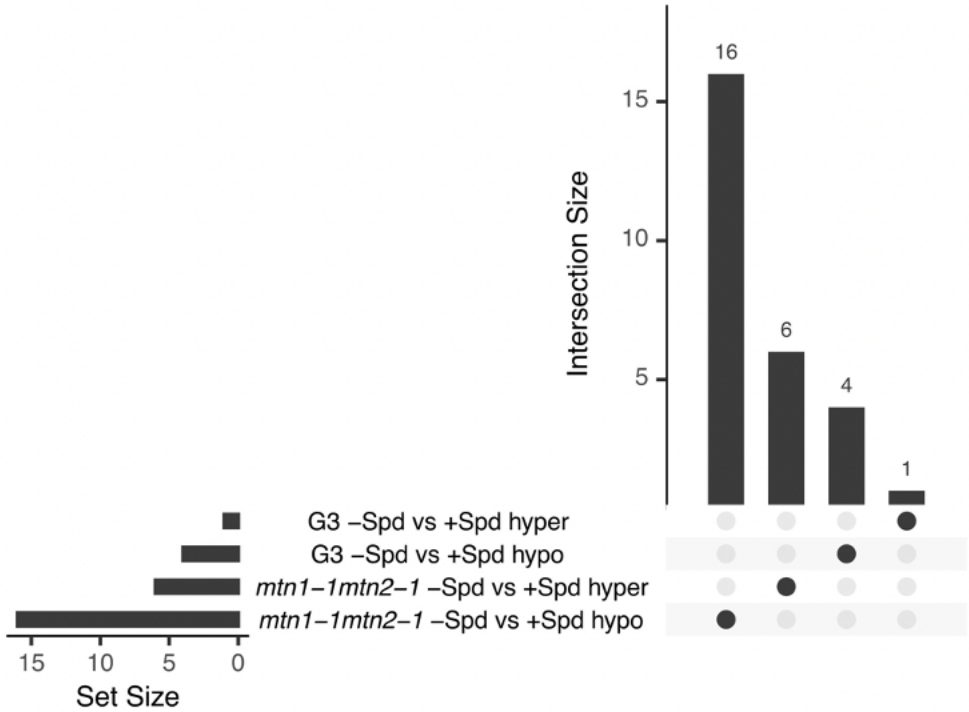


**b**

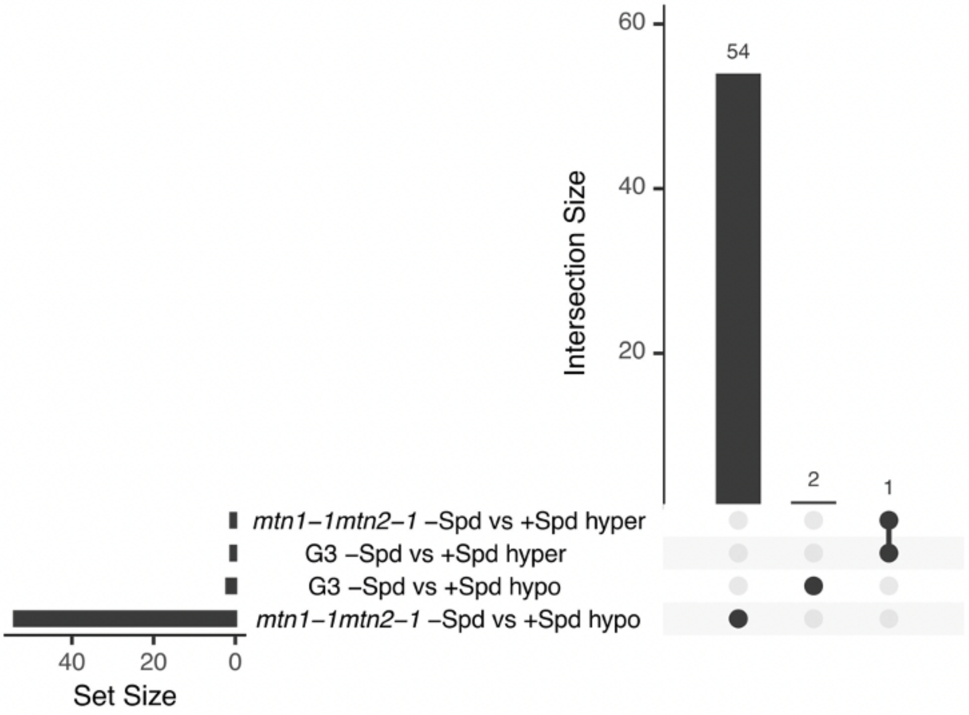


**Supplementary Figure 10: The effect of within-generation Spd supplementation on DNA methylation is minor.**

**a** UpSet plot of overlapping mCG DMRs in comparisons between BS-seq of Col-0 (WT), *mtn1-1mtn2-1*, and *mtn1-1mtn2-5* buds without Spd supplementation. **b** UpSet plot of overlapping mCHG DMRs from comparisons between treatments (with and without Spd supplementation) of samples from **a**.
