## Supplementary Note 1 for "Impaired methyl recycling induces substantial shifts in sulfur utilization in Arabidopsis"

**Supplementary Note 1: Detection of increased secondary sulphur assimilation in *mtn1-1mtn2-1.***

In the course of our HPLC measurements of SAM, SAH, and MTA in WT and *mtn1-1mtn2-1* buds, we noticed additional peaks in the resulting chromatograms with significantly higher signal in the mutant sample (Supplementary Note 1 Figure 1a). We suspected these peaks could be indicative of increased levels of adenosine-containing compounds such as APS, PAPS, or PAP as a result of increased secondary sulfur assimilation (Figure 2a), though we lacked the necessary standards to test this theory. As an alternative, we performed LC/MS to qualitatively compare any WT and *mtn1-1mtn2-1* LC/MS peaks with expected mass (m/z) and retention patterns indicative of these compounds. Using ion trap negative mode, we detected a peak in the *mtn1-1mtn2-1* sample with a mass of 507.9234 m/z, absent from WT, possibly matching PAPS (Supplementary Note 1 Figure 1b). With ion trap positive mode, we detected peaks with mass 287.0874 m/z and 355.1719 m/z in *mtn1-1mtn2-1*, absent in WT, matching the expected mass of adenosine and APS, respectively (Supplementary Note 1 Figure 1c). Two additional peaks with mass 433.1733 m/z and 595.2594 m/z, matching the expected values of PAP and PAPS, respectively, could be detected in both samples. From these results we hypothesized that normal levels of sulphate in *mtn1-1mtn2-1*, in combination with reduced primary sulphur assimilation, is leading to increased APS and PAPS, indicative of increased secondary sulphur assimilation. Additional work will be required to validate these data.

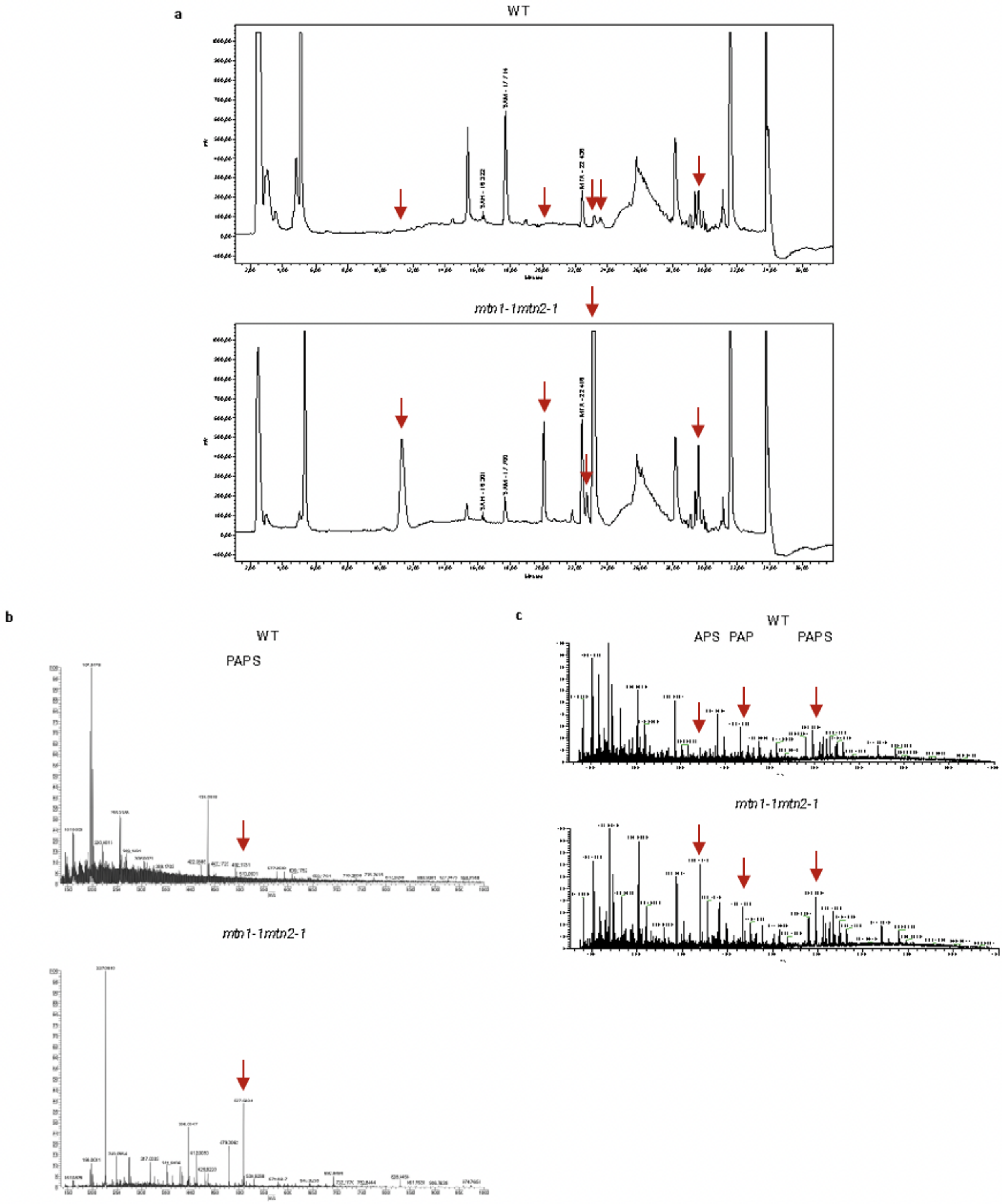


**Supplementary Note 1 Figure 1: Unknown peaks in *mtn1-1mtn2-1* HPLC may be indicative of increased APS and PAPS in *mtn1-1mtn2-1*.**

**a** Representative HPLC chromatogram from WT and *mtn1-1mtn2-1* buds samples run on a Nova-Pak C18 column to quantify adenosine-containing compounds SAM, SAH, and MTA. Arrows point to unknown peaks with increased signal in the *mtn1-1mtn2-1* samples. **b** LC/MS traces of mass (m/z) of compounds detected in WT and *mtn1-*

*1mtn2-1* bud samples using ion trap negative mode with a Velos LTQ Orbitrap Pro (ThermoFisher Scientific). Arrows point to the putative PAPS peak with mass 507.9234 m/z. **c** Same procedure as **b** using ion trap positive mode. Arrows point to putative adenosine, APS, PAP, and PAPS peaks with mass 287.0874 m/z, 355.1719 m/z, 433.1733 m/z and 595.2594 m/z, respectively.
